# Morphological parameters can capture emergent properties of dynamic disordered cytoskeletal networks

**DOI:** 10.64898/2026.03.01.708800

**Authors:** Soumik Ghosh, Lilianna Houston, Alejandra Vasquez, Kingshuk Ghosh, Ashok Prasad

**Affiliations:** School of Biomedical and Chemical Engineering, Colorado State University, Fort Collins, Colorado, USA; Department of Physics and Astronomy, University of Denver, Denver, Colorado, USA

## Abstract

The actin cytoskeleton is an inherently disordered active system. the actomyosin cortex and reconstituted actomyosin systems are globally disordered, yet undergo transitions between distinct disordered states as parameters like motor and crosslinker concentration and filament length and rigidity change. In cells these changes are related to genetic mutations or differences in cell state and dictate fundamental biological processes. However, we don’t have well established methods to detect and classify differences in disordered polymer networks. Image-based morphology techniques provide a non-invasive, high-throughput method of extracting information about a system. In this work we simulate biopolymer networks under varying conditions and develop and use morphological descriptors to construct trajectories in morphospace. Using statistical analysis we find that morphological descriptors are able to distinguish between different trajectories of the system, including differences not apparent to the eye. However, no single descriptor alone is able to capture all the differences in the simulated trajectories. Nematic order parameters typically perform the worst for our simulations while curvature and texture descriptors can collectively distinguish between dynamic trajectories. This work helps develop quantification of cytoskeleton dynamics for classification and data-driven modeling.

## 1 Introduction

The actin cytoskeleton is one of the fundamental determinants of cell function, serving as both a structural scaffold and a dynamic engine for mechanical processes. It is being increasingly recognized that the specific architecture of cellular actin determines how a cell grows, moves, generates force, and responds to its environment[1, 2, 4, 9, 10, 16, 20, 24, 25, 26, 27]. Around 400 proteins are believed to control actin architecture, from polymerization to higher-order networks[11] underlining its importance in cell function. In the generation of force for example, cells transition between different actin architectures to modulate their contractile output [5, 24, 25, 26]. In platelets, radial versus circumferential filament orientation determines the magnitude and localization of traction forces, critical for stable adhesion during the clotting process [20]. Similarly, cytoskeletal architecture is inextricably linked with function for the other cytoskeletal networks (microtubules and intermediate filaments) in cells.

While there are a few important phenomena in which the cytoskeleton is ordered such as the actin contractile ring in mitosis[19], the microtubule spindle during interphase, or actomyosin sarcomeres in muscle cells, in most cases the cytoskeleton presents itself as a disordered dynamic network. Nevertheless, identifiable attractor states in response to external perturbation[9] or genetic mutations[2] have been observed, suggesting that there are yet to be discovered higher-order principles that govern the cytoskeletal architecture. The significance of cytoskeletal architecture raises the question of how it can be described quantitatively. Since much of the information about network organization derives from light microscopy, a natural framework for describing cytoskeletal organization is through image-derived morphological metrics. Morphological metrics have been used to classify static images of the actin network that differ by chemical or genetic perturbations [9, 1, 2], but we do not know how useful they are in distinguishing between dynamic trajectories of the network that may differ due to differences in the underlying cell states. Being able to capture and quantify these differences is essential for comparing experiments across various conditions and for linking image-based morphology to the mechanics of active matter. This creates the need for identifying a set of morphological parameters that can distinguish the various states of such disordered systems. In this paper, for reasons discussed below, we chose to utilize measures of filament curvature and image texture.

Cytoskeletal polymers are semiflexible, i.e., they have bending rigidity characterized by a persistence length which ranges from around 20*µm* for actin to millimeters for microtubules when measured *in vitro* [12]. Despite their high bending rigidity microtubules are frequently curved in cells due to the compressive loads they bear from the contractile actomyosin cytoskeleton[8]. The intermediate filament vimentin is also highly curved and even wavy in cell images [28] and forms rings around the nucleus during cell spreading [29]. Actin typically assembles into shorter linear filaments that form branched networks, however, force-induced curvature of actin filaments has been observed[23]. Specialized proteins can also curve actin filaments, like septins or IQGAP (IQ-motif containing GTP-ase proteins) [21]. Since curvature encodes substantial information about force generation and transmission within the cell and its surroundings, we constructed curvature-based measures, described in detail in the Methods section, to capture these effects.

Of the components that make up the actin cytoskeleton, the actin cortex and the lamellipodia are highly cross-linked networks of predominantly branched linear filaments. Differences in network organization to eye resemble differences in texture, motivating the application of textural measures to characterize the network structure.

Because cytoskeletal and cellular systems are inherently disordered and spatially heterogeneous, Haralick features are particularly well-suited for capturing emergent structural patterns that arise from collective filament organization and active processes. Haralick features are one of the most widely used texture descriptors, computed from the gray-level co-occurrence matrix (GLCM), which summarizes how often intensity value pairs occur for a given distance and direction. The GLCM data is then converted into interpretable measures, such as contrast, correlation, energy, and homogeneity [13]. Haralick feature analysis has been used on a wide variety of cytoskeletal systems. They have been shown to be informative in classifying cell types [3], quantify changes in actin architecture during wound repair [15]), measure changes in microtubule organization [17, 7] and vimentin networks [31].

We reasoned that using both curvature and texture measures would allows us to capture complementary information about system state. Curvature encodes the effect of forces at the filament level, while the texture features encode network organization of a different type. We then asked whether these morphological metrics can distinguish between differences in polymer network dynamics due to differences in basic parameter values.

We address this question using simulations, where individual parameters governing assembly like filament length, rigidity, motor and crosslinker concentrations, can be systematically varied. This allows us to directly assess the sensitivity of curvature and texture features to specific model parameters.

## 2 Materials and Methods

### 2.1 System

#### 2.1.1 Simulation System

We simulate a polymer network using Cytosim [22], containing filaments, motors and crosslinkers. We adapted and developed a previously published simulation [6, 32] on the actin cytoskeleton. In contrast to earlier work, we intentionally sample a broader and more diverse parameter space, generating a wide set of systems with distinct combinations of biophysical parameters, that can be explored using our morphological measures. All simulations are carried out in 2D for simplicity and computational speed. Each simulation runs for 200000 frames with a timestep of 0.001*s* each, from which 600 evenly spaced frames were recorded.

By varying motor concentrations and filament length, we observe four distinct phenotypes:

1. Low-motor / small-filament (LM–SF): weakly organized networks with slight contractility.
2. High-motor / small-filament (HM–SF): strong condensation into compact aggregates or asters.
3. Low-motor / large-filament (LM–LF): largely unorganised networks with minimal contractility.
4. High-motor / large-filament (HM–LF): condensation into aggregates or asters that are less compact than in the small-filament case.

Motor densities are set to 6.25 and 100 motors per µm*^−^*^2^ for the low-motor (LM) and high-motor (HM) regimes, respectively, while the crosslinker density is initially fixed at 250 per µm*^−^*^2^. The total filament length across all filaments is held constant across all simulations. In large-filament (LF) systems, the average filament length is approximately 1.54 µm. Small-filament (SF) systems are generated by splitting each large filament into two equal segments.

Although all four states are globally disordered, their local morphology encodes distinct biophysical parameters of the system. Low-motor (LM) systems exhibit a stalled network, whereas high-motor (HM) systems undergo condensation that is readily discernible by eye. While systems with different filament lengths behave similarly at the same motor concentrations, subtle differences arise in these cases, that are not readily apparent by visual inspection.

We also sample the previous cases over two rigidities: 0.075 pN µm*^−^*^1^, which is the same as actin filaments (AR) which should be more easier to bundle, and 0.01 pN µm*^−^*^1^, which reflects a filament of much lower rigidity (LR), which should be much more pliable to the motors. This interplay should result in systems that show similar general trends but are still distinctly different.

Because crosslinker-mediated stabilization and bundling play a crucial role in determining the final system state at late trajectory times ([6]), so, in addition to the initial “high” regime (HCL, density=250) used previously, where bundling should predominate at late trajectory times, we also sample a “low” regime (LCL, density=15.625), where this crosslinker mediated stabilization is significantly reduced.

Simulations take place in a box of side 4 µm. Since in real life, filaments will likely stay in confined systems, we examine the role of external space constraints. We perform an additional set of simulations in which the cytoskeletal network is confined within a fixed circular boundary representing a simplified cell-like circular geometry of diameter 4 µm. This constraint introduces geometric and mechanical restrictions that should alter filament organization, allowing us to directly assess how confinement modifies the curvature and texture metrics relative to the unconfined systems.

Across this parameter space, we aim to use different morphological descriptors to distinguish between and gain insights about systems that may appear qualitatively similar, but arise from distinct underlying biophysical regimes.

#### 2.1.2 External Validation on Published Cytoskeletal Images

To validate our proposed morphological parameters, we also apply a subset of them to a cell images published by Hauke et al. [14]. All experimental procedures, including staining of cytoskeletal filaments (actin,vimentin,microtules), substrate preparation across five stiffness conditions, and image acquisition, were carried out in the original study. Here, we use only the published images as an external validation dataset.

### 2.2 Morphological parameters

We look at 3 scales of parameters for our study.

1. Global Textural parameters: Haralick features
2. Filament level features: Nematic order parameters, Mean alignment order of filaments
3. Local features: Mean curvature of filament segments

We explore the behavior of these features across an ensemble of 20 simulations each for all the cases mentioned above.

#### 2.2.1 Haralick Features

Haralick texture features are statistical descriptors derived from the gray-level co-occurrence matrix (GLCM), originally introduced by Robert Haralick [13] to quantify spatial relationships between pixel intensities in images. These features capture key aspects of image texture such as contrast, homogeneity, entropy, correlation, and energy, providing a compact numerical representation of structural organization beyond simple intensity-based metrics.

In our work, we use Haralick features to serve as quantitative texture descriptors to characterize spatial organization within cytoskeletal networks, enabling systematic comparison across experimental conditions without worrying about segmentation or filament tracking.

Haralick features are sensitive to free space. For experimental images which are not as sharply resolved as our simulation images, segmenting the cell allows the calculation of the Haralick features as normal, ignoring the transitions to and from 0 intensities in the GLCM. This makes it consider only the object system and not the empty space around it. Since our simulation results have such high resolution and our non-filament background is 0, ignoring the zero transitions would lead to calculation of texture only along the filaments. If we do not ignore the zero transitions, it would lead to classification of systems that do and do not condense, based mainly on the free space, instead of internal texture due to the filament system. So we try to find a reasonable area that contains a representative snapshot of the system, without too much free space. We do this by asymmetrically cropping these systems around the centroid of the system, such that the cropped region contains 70% of the bright pixels, before calculating Haralick features on the resulting simulation images.

#### 2.2.2 Nematic Order Parameters

The nematic order parameters are a measure of how aligned a collection of elongated objects is. They are commonly used to quantify alignment in actin bundles, stress fibers, and contractile structures, and are especially powerful for studying order–disorder transitions. We test four of the morphological parameters laid out by Akenuwa et al. ([1]), namely, Mean Angle([18]), Angular Variation, Parallelness ([30]) and Order Parameter. Mean Angle is a measure of orientation while the other three are a measure of ordering. But for our active disordered systems, they do not perform well (Figure SI1), probably because our simulation systems do not show any nematic ordering.

#### 2.2.3 Mean Curvature of Filament Segments

Mean curvature is a measure of the average degree of filament bending within the network. Filament coordinates obtained from Cytosim are discretized into segments of length 0.5 µm. For each filament, curvature is computed at each segment using the standard planar curvature expression

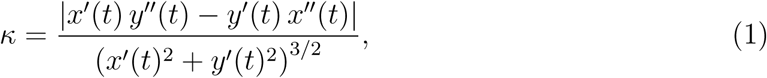

where *x*(*t*) and *y*(*t*) denote the cartesian coordinates of a filament and the primes indicate derivatives taken along the filament. The numerator measures how rapidly the filament changes direction, while the denominator normalizes this change to yield a geometric measure of curvature.

The absolute value of *κ* is retained so that curvature reflects bending magnitude independent of direction. Curvature values are computed for all filament segments and the overall mean for the system is retained.

#### 2.2.4 Mean Alignment Order Of Filaments

The mean alignment order of filaments is a measure of the average curvedness of filaments of a system, compared to a system with only straight filaments. For each filament, we create a “director”, which is a straight line joining the ends of a given filament. We then take each filament section of size 0.5 µm as discussed earlier, and calculate the angle each section makes with its respective filament director using dot product. Let **v***_i_* = (Δ*x_i_,* Δ*y_i_*) denote the local segment direction, and let **v**_dir_ = (Δ*x*_dir_, Δ*y*_dir_) denote the filament director. The angle *θ_i_*between the segment and the director is then given by

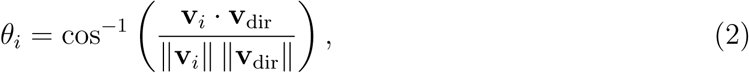

where **v***_i_ ·* **v**_dir_ is the dot product and ││·││ denotes the Euclidean norm. Angles are computed for all filament segments and and the overall mean for the system is retained.

### 2.3 Principal Component Analysis (PCA)

Principal Component Analysis (PCA) provides an low-dimensional representation of the data that preserves the most variance along orthogonal directions. It is used to reduce the dimensionality of the feature space and to summarize dominant linear patterns in the data. PCA was applied to our standardized filament-level and Haralick texture features.

PCA was computed using a singular value decomposition–based algorithm, calculating the top 10 principal components for each observation and retained along with metadata identifying experimental condition, time frame, and replicate.

The resulting principal component scores were used for visualization and exploratory analysis. The first 2 components explained a lot of the variance, so they were plotted for visualization, along with time-resolved visualizations to track the evolution of observations in principal component space. PCA loadings and explained variance ratios were examined to assess the contribution of individual features to each component and to guide interpretation of dominant linear patterns in the data.

### 2.4 Uniform Manifold Approximation and Projection (UMAP)

To complement PCA and possibly decode nonlinear structure in the data, Uniform Manifold Approximation and Projection (UMAP) was employed as a nonlinear dimensionality reduction technique for visualization. UMAP was applied to the same standardized feature sets used in the PCA analysis.

UMAP was computed by first creating a weighted nearest-neighbor graph in the original high-dimensional feature space, from which a low-dimensional embedding is created that optimally tries to preserve local neighborhood relationships while maintaining aspects of the global structure. UMAP embeddings were calculated using a Euclidean distance metric on standardized features, with the number of neighbors=30 and the minimum distance parameter=0.05. We sampled different values for the number of neighbours (15,30,50) that have been used in literature and provided the others in supplementary information (Figures SI2 to SI9). We decided to use 30 as it seems to decode local feature clustering well enough, without losing global trends.

The UMAP embedding was fit once globally using all observations, and the resulting low-dimensional coordinates were retained for visualization and temporal analysis. Observations were visualized in two-dimensional UMAP space, colored by experimental condition, and time-resolved visualizations were constructed by slicing the global embedding by frame to track the evolution of system dynamics over time.

To aid interpret the embeddings, Pearson correlations between original standardized features and UMAP coordinates were also computed.

## 3 Results

### 3.1 Simulation data reveal differences in dynamic trajectories in different parameter regimes

Representative snapshots of our simulations are shown in Figure 1 for a box geometry of size 4*µm*. Panel I has snapshots corresponding to filaments with actin’s rigidity (AR) and a high cross-linker concentration (HCL). These images show significant differences in the dynamics for the low motor (LM) concentrations (A & C) versus the high motor (HM) concentrations (B & D), with the HM simulations showing condensation to a central aster-like condensed structure at early times while the LM simulations show a stalled network. For the HM cases, there is also a clear difference between the short filaments (SF) in row B versus the longer filaments (LF) of row D, where the former shows only partial condensation as compared with the latter. Less easily apparent to the eye, the SF (row A) and LF (row B) cases with the LM concentrations show differences in bundling with time. We should reiterate here that all simulations start from the same initial conditions (see *Materials and Methods*).

**Figure 1:**
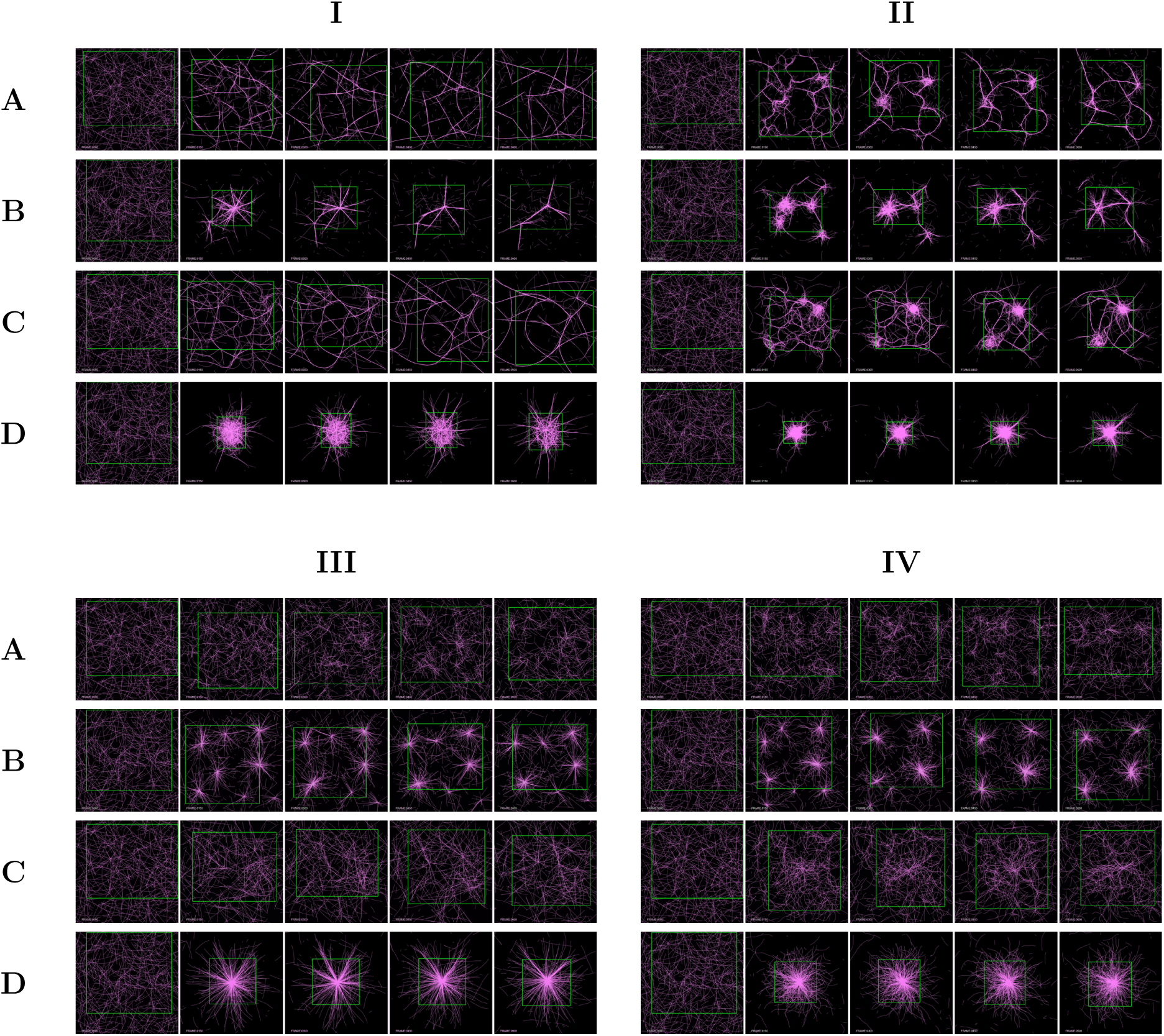
System evolution of filament trajectories in a box of side=4. Panels I-IV correspond to AR-HCL, LR-HCL, AR-LCL, LR-LCL. Rigidities: AR = 0.075 pN µm*^−^*^1^, LR = 0.01 pN µm*^−^*^1^. Crosslinker densities: HCL = 250, LCL = 15.625. Rows: *(top to bottom)* A=LM-SF, B=HM-SF, C=LM-LF, D=HM-LF. Columns: *(left to right)* frames=0, 150, 300, 450, 600.

Panels I-IV are organized similarly. In Panel II we see that decreasing the rigidity of the polymer (LR) at HCL increases the ability of motors to condense filaments. More enhanced bundling can be seen in row A (SF-LM), and aster-like structures form in all the other cases, including the LM-LF case (row C). Panels III and IV show what happens when we drop cross-linker concentrations (LCL) with either actin rigidities (panel III) or low rigidities (panel IV). While row A (LM-SF) in both cases looks generally unaffected, a higher motor concentration (row B) leads to multiple asters forming (row B). Interestingly, the lower cross-linker concentration is insufficient to form condensed structures when the motor concentration is low for the longer filaments. Row D looks similar across all panels.

In Figure 2 we show representative snapshots from simulations in a circular box of diameter 4*µm* which acts as a small geometric constraint compared with the box. The effect of this constraint is to enhance condensation, which we can see by comparing the global organization of condensation in Figure 2, Panel I and II with its counterpart for the square cell. Similarly, constraining the filaments to a circular cell (Figure 2, Panel III and IV) in the “low” crosslinker regime, helps to achieve global condensation again.

**Figure 2:**
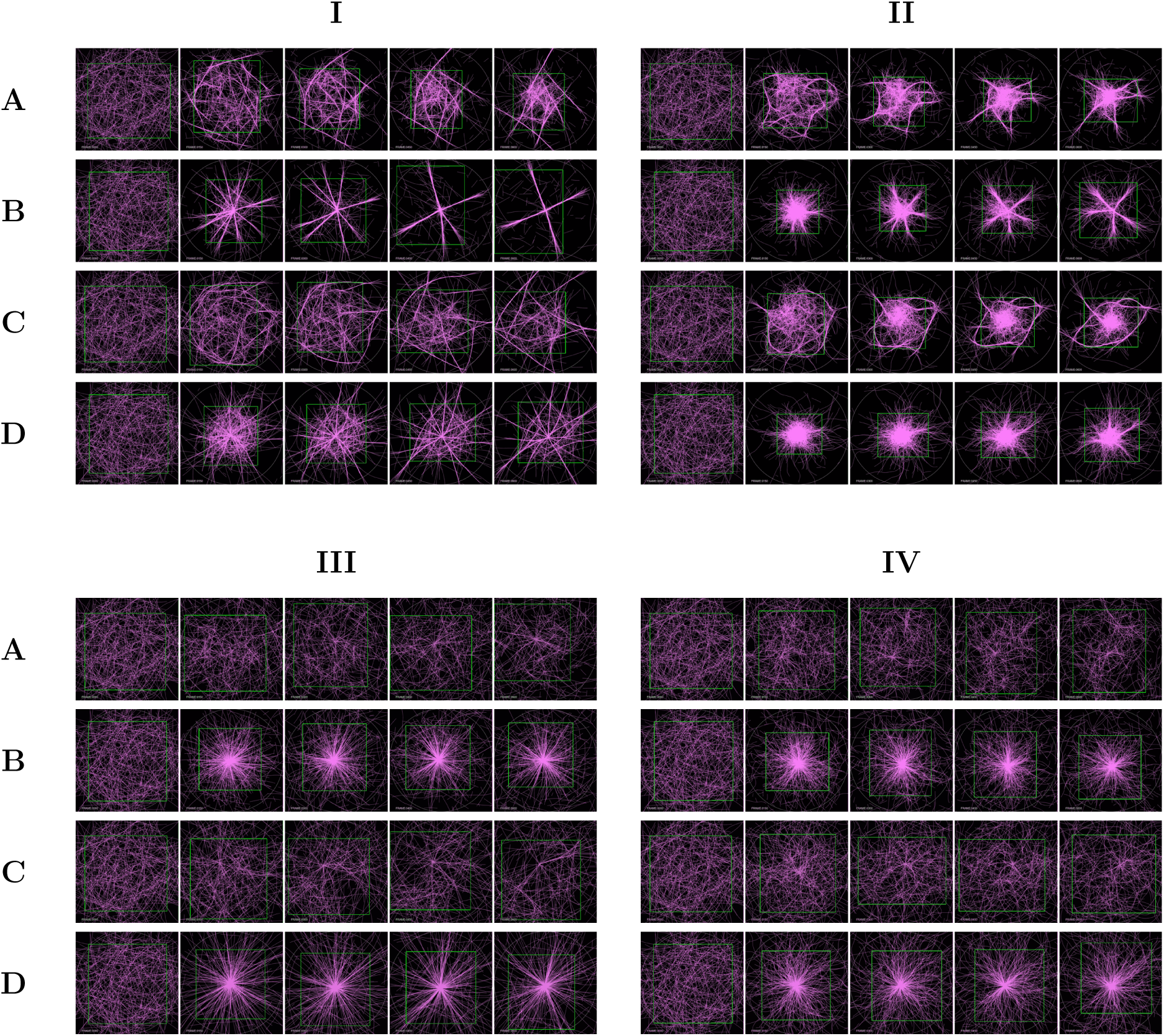
System evolution of filament trajectories in a constrained circular cell of diameter=4. Panels I-IV correspond to AR-HCL, LR-HCL, AR-LCL, LR-LCL. Rigidities: AR = 0.075 pN µm*^−^*^1^, LR = 0.01 pN µm*^−^*^1^. Crosslinker densities: HCL = 250, LCL = 15.625. Rows: *(top to bottom)* A=LM-SF, B=HM-SF, C=LM-LF, D=HM-LF. Columns: *(left to right)* frames=0, 150, 300, 450, 600.

#### 3.1.1 Filament-level features show separation based on mechanical factors

Filament-level descriptors are based on polymer bending, either represented as the angular difference from the end-to-end vector or as local curvature (see Materials and Methods). Since polymer bending depends upon rigidity as well as segment length, these measures report on polymer mechanics and local geometry, and their temporal evolution reveals how motor concentration, filament length, rigidity, crosslinker density and confinement shape overall network organization. For all parameter regimes, both filament-level features show the most temporal variation during a early-trajectory “condensation phase” (from frame 0 to around frame 75, changing for different cases), followed by gradual convergence as the system transitions into a “bundling phase” at later trajectories. (Figure 3 and Figure 4).

**Figure 3:**
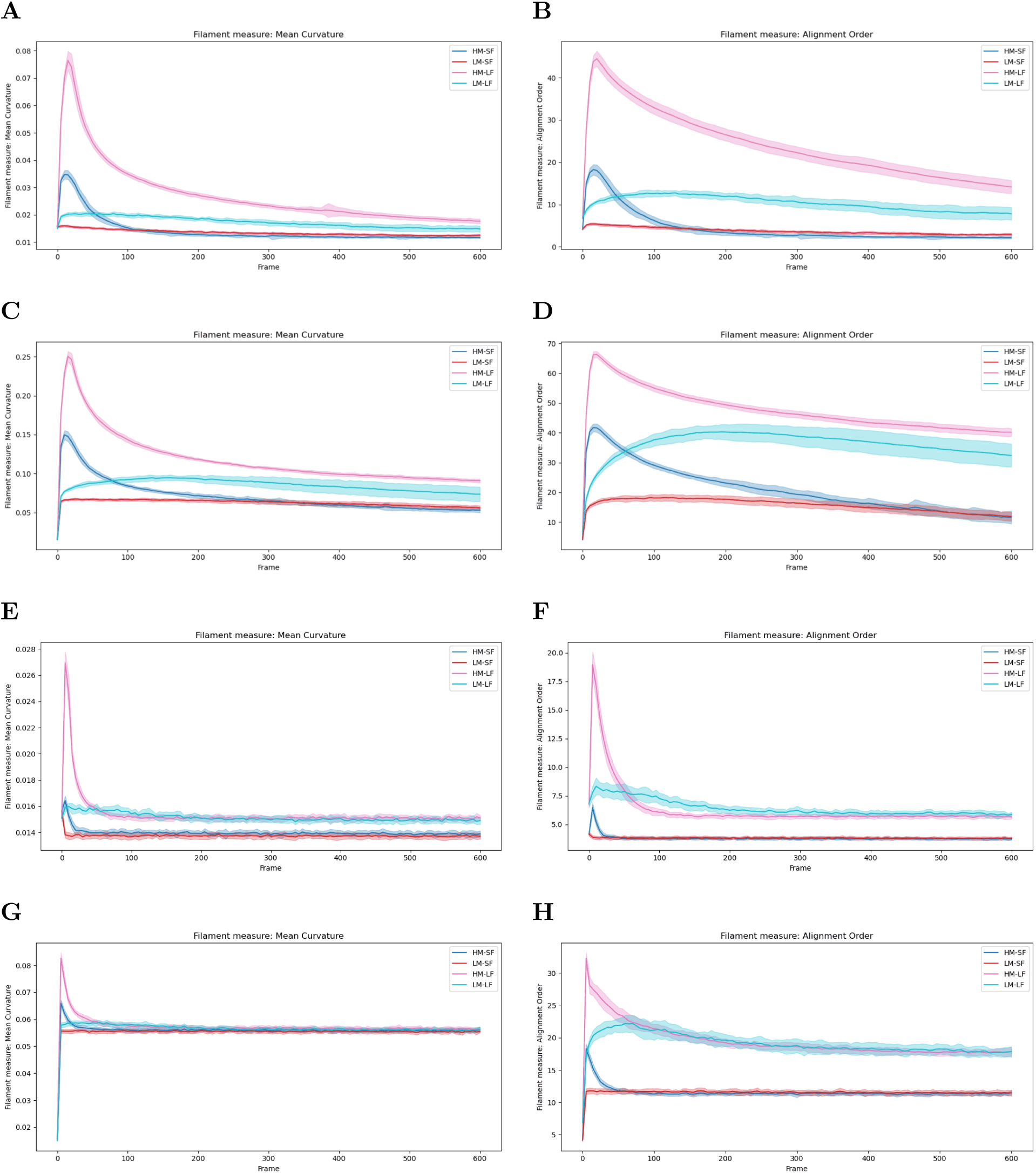
Ensemble averages of the evolution of filament features for box of side=4. Mean curvature (A,C,E,G) and alignment order (B,D,F,H) for rigidities 0.075 pN µm*^−^*^1^ (A,B,E,F) and 0.01 pN µm*^−^*^1^ (C,D,G,H), and crosslinker densities 250 (A-D) and 15.625 (E-H).

**Figure 4:**
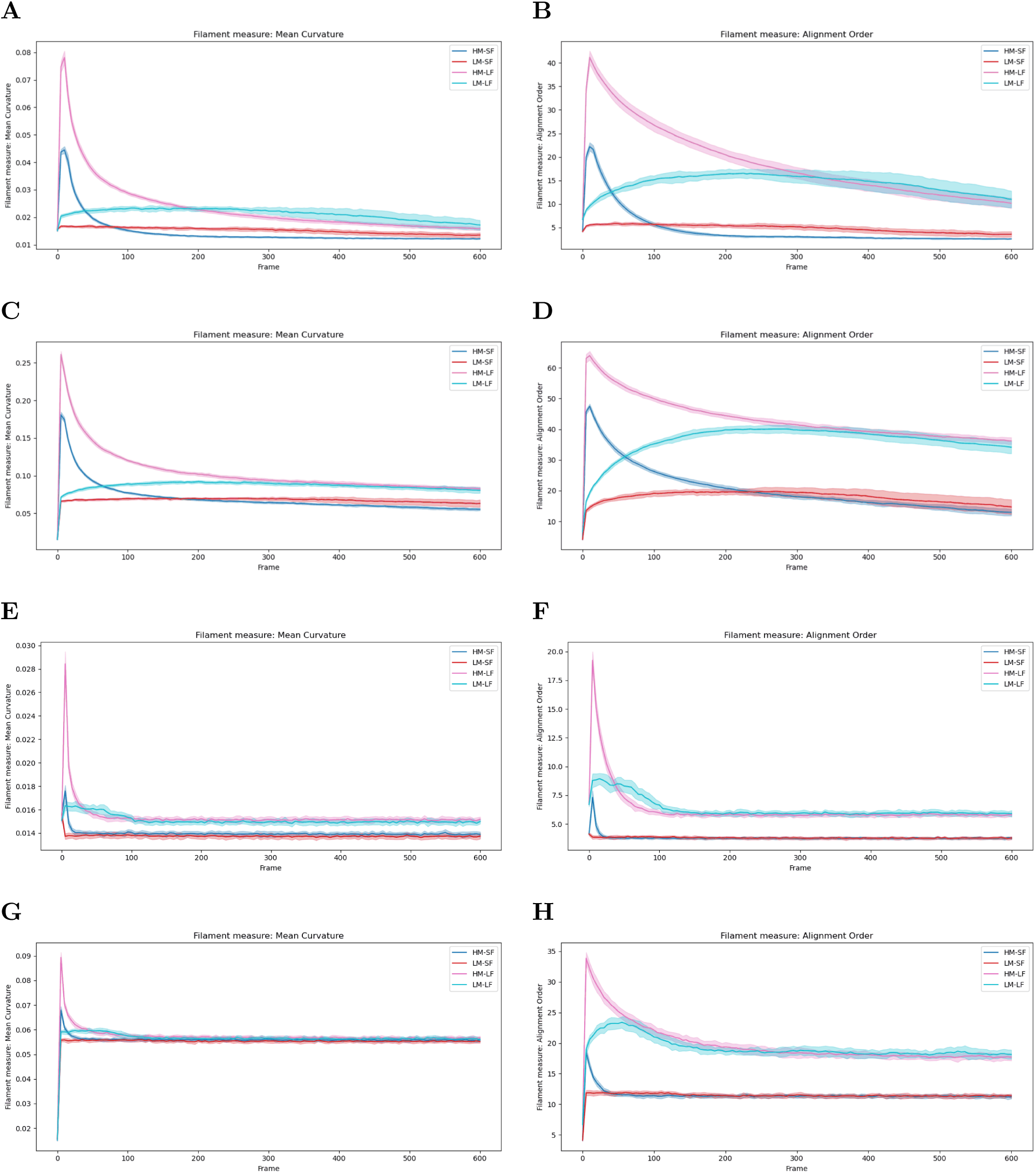
Ensemble averages of the evolution of filament features for a constrained circular cell of diameter=4. Mean curvature (A,C,E,G) and alignment order (B,D,F,H) for rigidities 0.075 pN µm*^−^*^1^ (A,B,E,F) and 0.01 pN µm*^−^*^1^ (C,D,G,H), and crosslinker densities 250 (A-D) and 15.625 (E-H).

During condensation, motor-driven activity causes rearrangements that the network accommodates primarily through filament bending and reorientation. Curvature therefore increases sharply at early times, particularly in high-motor systems, as well as in low-rigidity filaments, where bending energy is lower. Geometric confinement further increases early curvature changes by probably increasing interactions between the filament and the boundary, forcing filaments to bend and reorganize due to the restricted volume.

Alignment order evolves during condensation as well, since motor-driven activity organizes filaments into local clusters and bundles. This rapid change probably reflects the system transitioning from an initially disordered configuration to a condensed state. The amount and speed of these changes depend on filament length and rigidity. More flexible filaments can bend more easily out of alignment, causing higher alignment order. Shorter filaments are easier to move with motors and might have more sharper changes in alignment, whereas longer filaments show a more gradual change due to motors.

As the system evolves, filament-level features begin to converge as it enters the bundling phase. With more crosslinker binding, filament-filament contact points are stabilised, rearranging filaments into mechanically coupled bundles. These bundles behave as stiffer composite structures, making them harder to bend and reducing curvature values. Since with the help of the stabilization of the crosslinkers, the filaments often “straighten” out, causing them to align much better with their directors, and causing a drop in alignment angles as well.

These trends show that filament-level features encode rich, regime-dependent information during condensation, but may lose discriminative power as filaments bundle and become harder to bend..

#### 3.1.2 PCA and UMAP of filament-level features cluster along the filament-length and filament-rigidity axes

We asked whether systems with different underlying parameters cluster distinctly in the space of morphological features. To examine this, we projected the high-dimensional feature space into lower dimensions using two commonly used methods: principal component analysis (PCA) and Uniform Manifold Approximation and Projection (UMAP).

Principal component analysis provides a compact representation of filament-feature evolution by projecting the bins of the angles for our filament-level features into a low-dimensional morphospace. We analyze each crosslinker density separately by performing PCA on systems that jointly vary motor density, filament length, and filament rigidity. We see that the first 2 PCAs account for more than 70% of the explained variance for all the cases (Figure 5), allowing us to plot them (Figure 6) against each other to gain insight about the systems. While the static PCAs show some separation, looking at the time-resolved PCA trajectories by explicitly tracking system evolution (Figure 8) offers significantly more insight.

**Figure 5:**
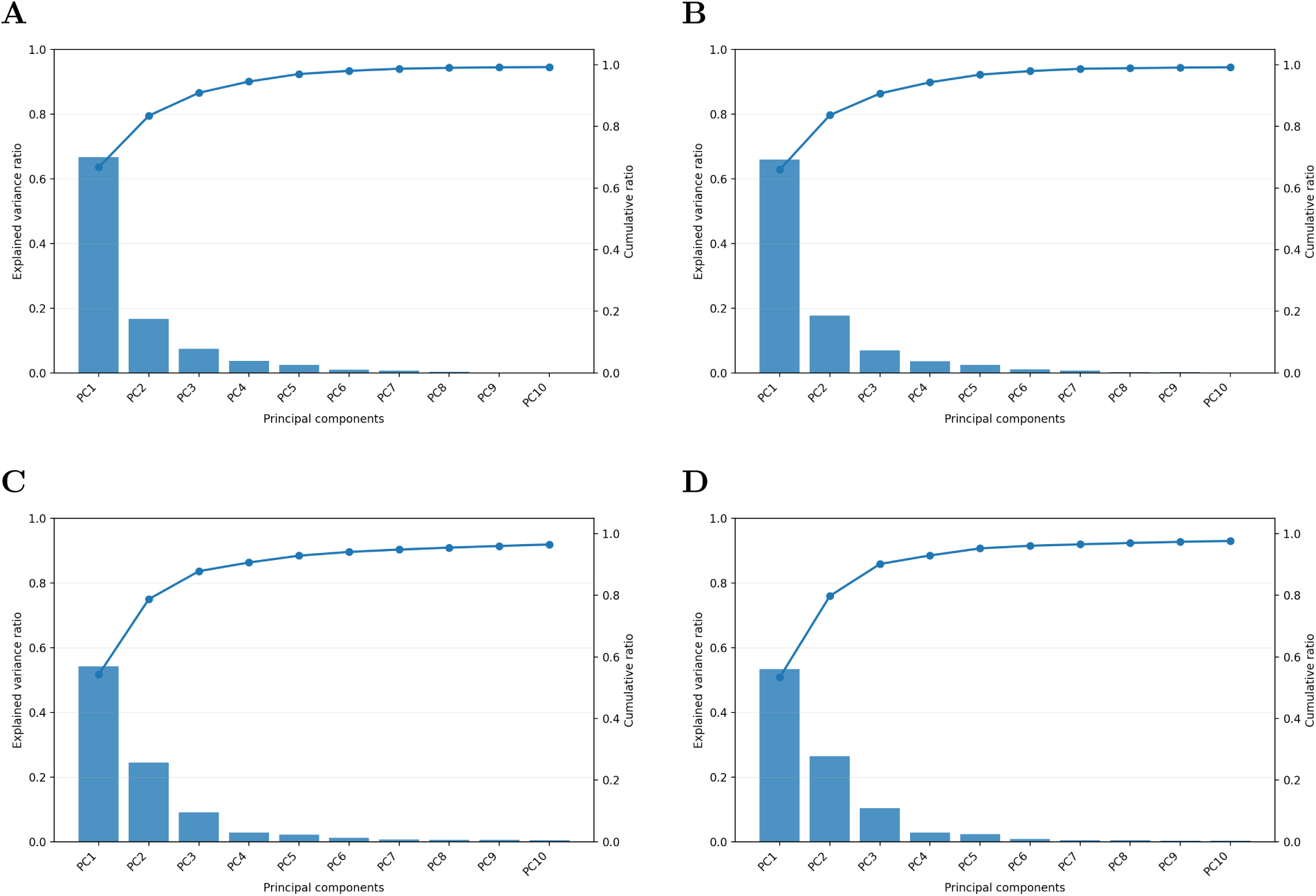
Explained variances for PCA plots for the bins of the filament features along with time for box of side=4 (A,C) and constrained cell of diameter=4 (B,D) including rigidities 0.075 pN µm*^−^*^1^ (AR) and 0.01 pN µm*^−^*^1^ (LR) for crosslinker densities 250 (A-B) and 15.625 (C-D).

**Figure 6:**
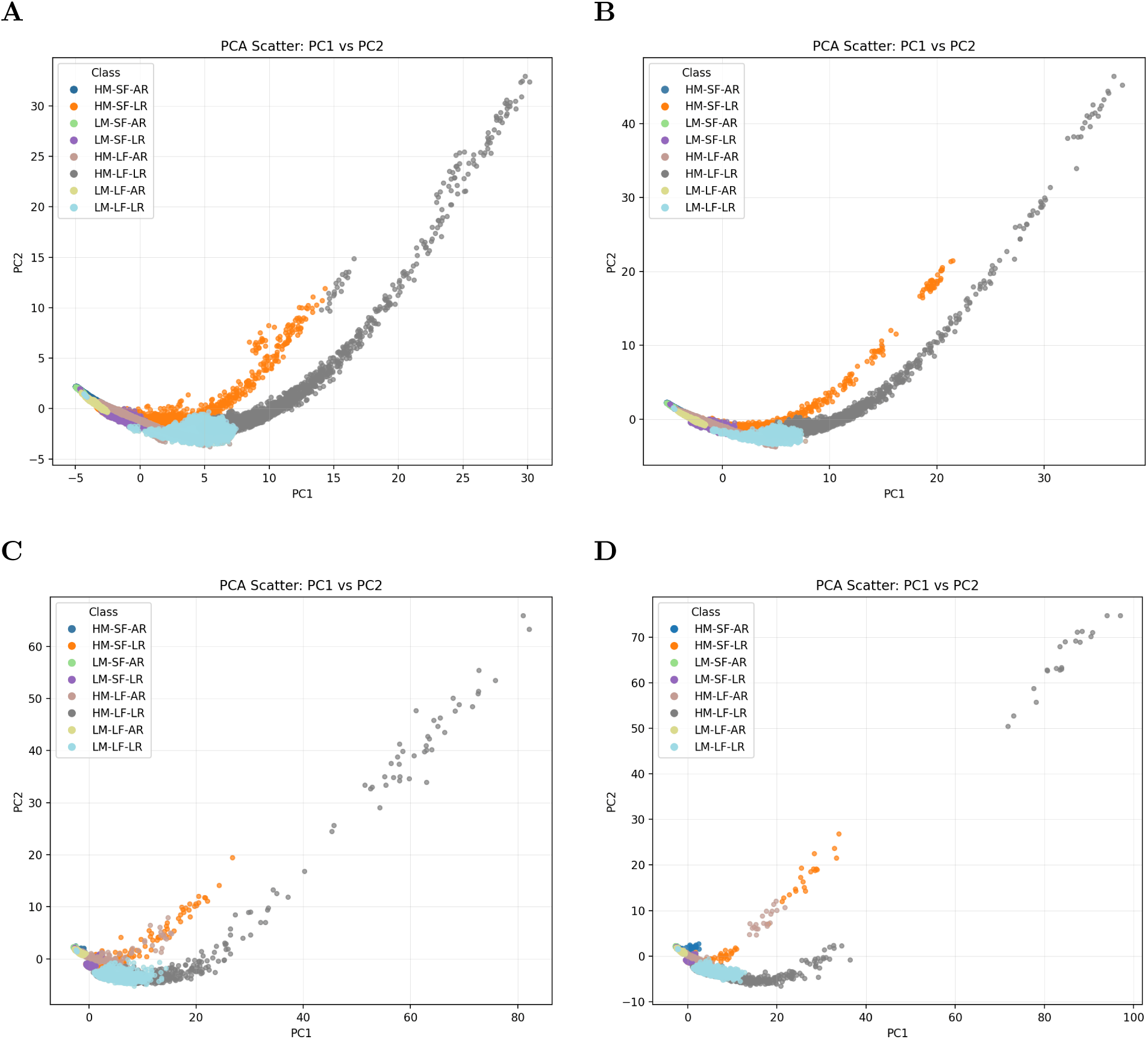
PCA plots for the bins of the filament features along with time for box of side=4 (A,C) and constrained cell of diameter=4 (B,D) including rigidities 0.075 pN µm*^−^*^1^ (AR) and 0.01 pN µm*^−^*^1^ (LR) for crosslinker densities 250 (A-B) and 15.625 (C-D).

After starting at the same location due to initial conditions, the systems typically diverge during condensation before converging back during bundling. Over time, the clusters mainly group themselves along the filament length-rigidity pairs. Low-rigidity pairs typically show more separation than high-rigidity pairs. At higher crosslinker density (Panel A-B), at long times, due to bundling, there is a loss of discrimination power and the higher rigidity groups slowly start to merge together. But as we go to lower crosslinker densities (Panel C and D), all short filaments group together, whereas the long filaments form two different groups based on rigidity.

Applying UMAP on the same standardized feature set as the PCA, offers a non-linear dimensionality reduction of the problem. For UMAP (Figure 7), like in PCA, we see that the cases with the same crosslinker density behave similarly to each other. But we see much better separation of the clusters. From the static UMAPs, we see that for the high crosslinker case (Panel A-B), only the short-filament cluster together with those of similar rigidity, but the different long filament cases cluster separately. But for the low crosslinker case (Panel C-D), we see clear groupings by filament-length and rigidity, forming 4 distinct groups.

**Figure 7:**
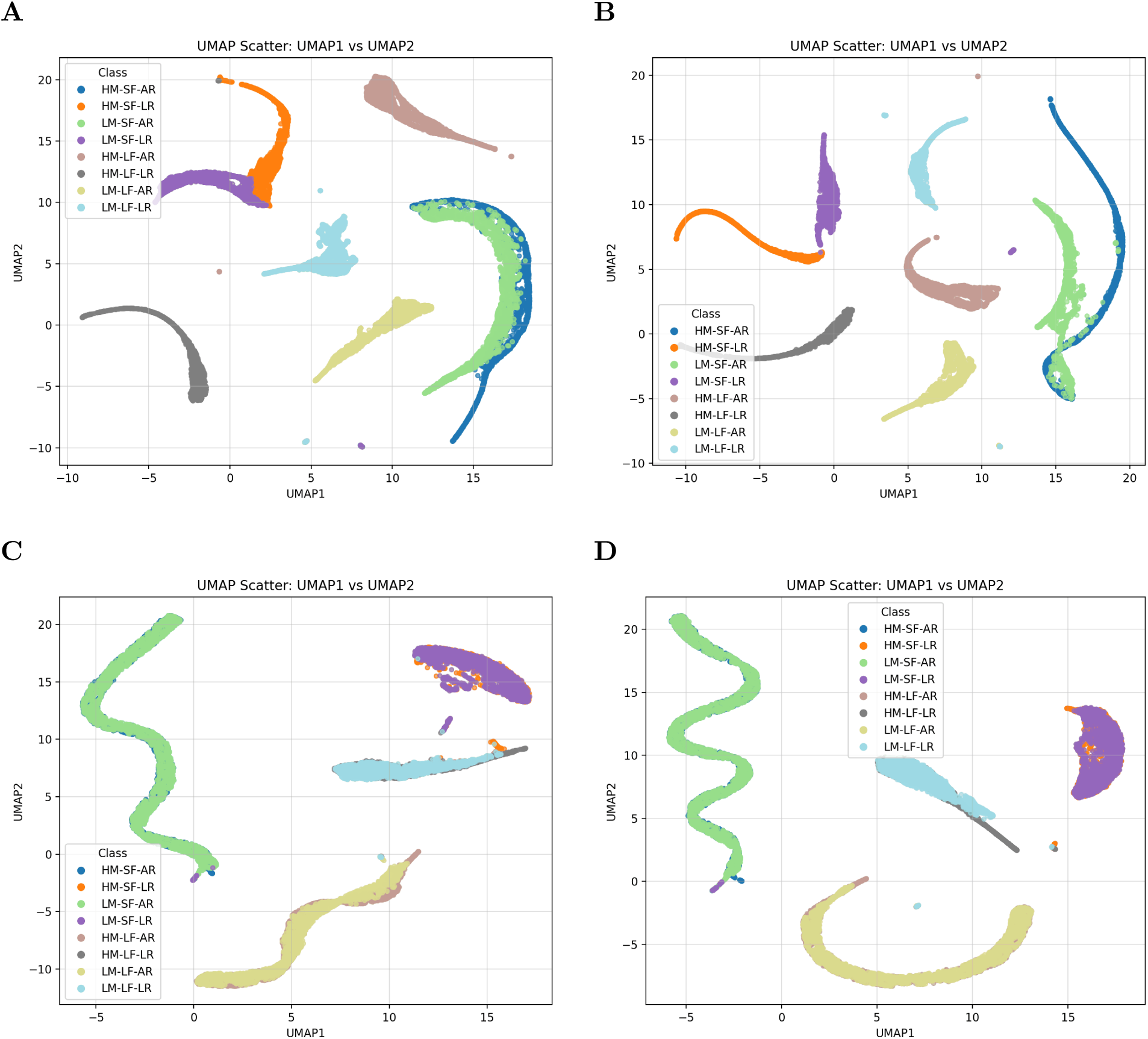
UMAP plots for the bins of the filament features along with time for box of side=4 (A,C) and constrained cell of diameter=4 (B,D) including rigidities 0.075 pN µm*^−^*^1^ (AR) and 0.01 pN µm*^−^*^1^ (LR) for crosslinker densities 250 (A-B) and 15.625 (C-D).

**Figure 8:**
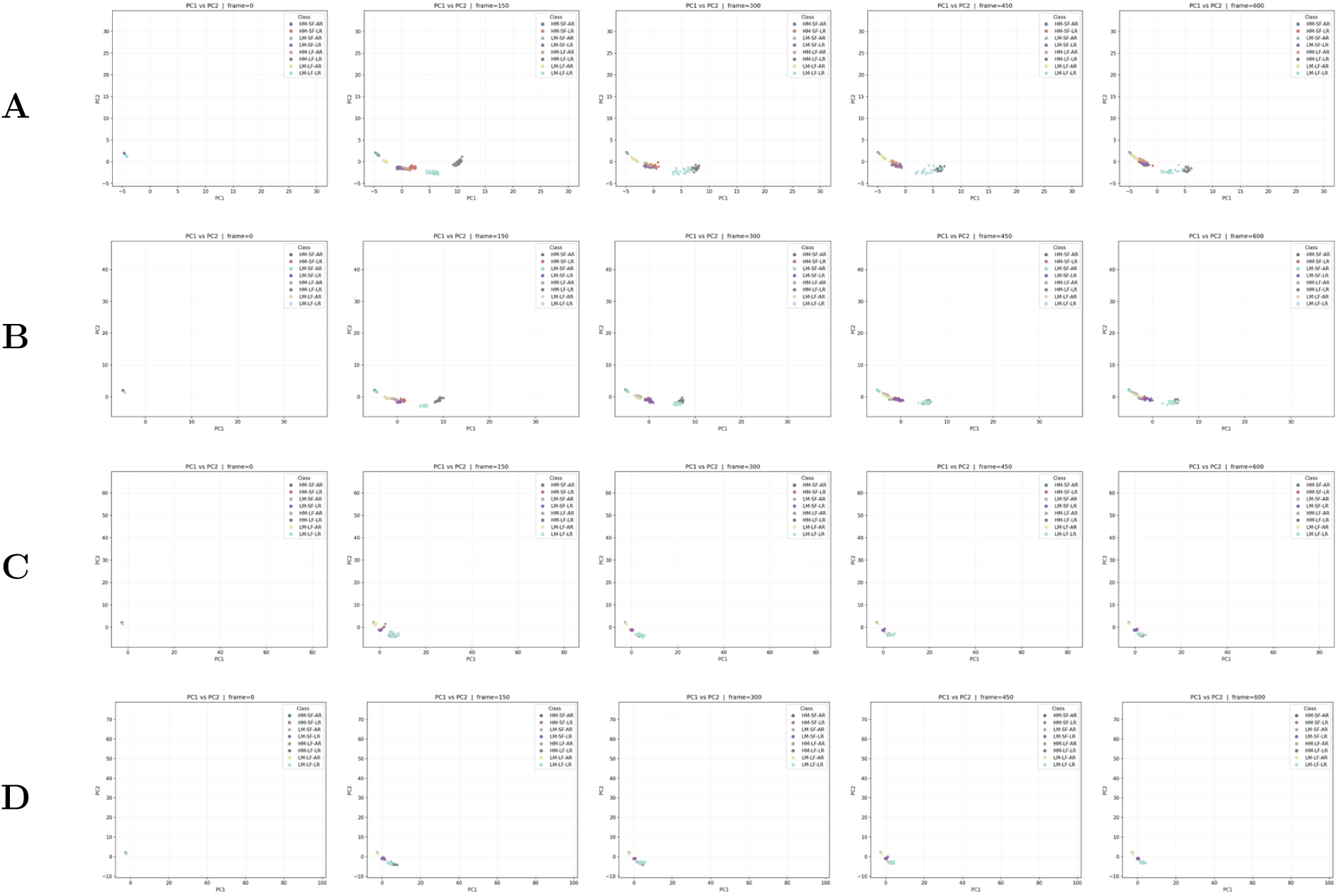
Evolution of trajectories of PCA plots for the bins of the filament features for box of side=4 (A,C) and constrained cell of diameter=4 (B,D) including rigidities 0.075 pN µm*^−^*^1^ (AR) and 0.01 pN µm*^−^*^1^ (LR) for crosslinker densities 250 (A–B) and 15.625 (C–D). Columns: *(left to right)* frames=0, 150, 300, 450, 600.

Checking the time-resolved UMAP trajectories (Figure 9), we see an initial separation from initial conditions, followed by convergence of the small filament groups with same rigidity for the high crosslinker cases. The extent of mixing for the small filament cases vary, with the small filaments of actin rigidity clustering as early as frame 150 for the box (Panel A) and as late as frame 450 for the constrained case (Panel B). The low-rigidity short-filament cases also converge but take longer than the higher rigidity cases. For the low-crosslinker case (Panel C-D), the 4 groups for filament length and crosslinkers form early after initial divergence and remain grouped throughout the trajectory.

**Figure 9:**
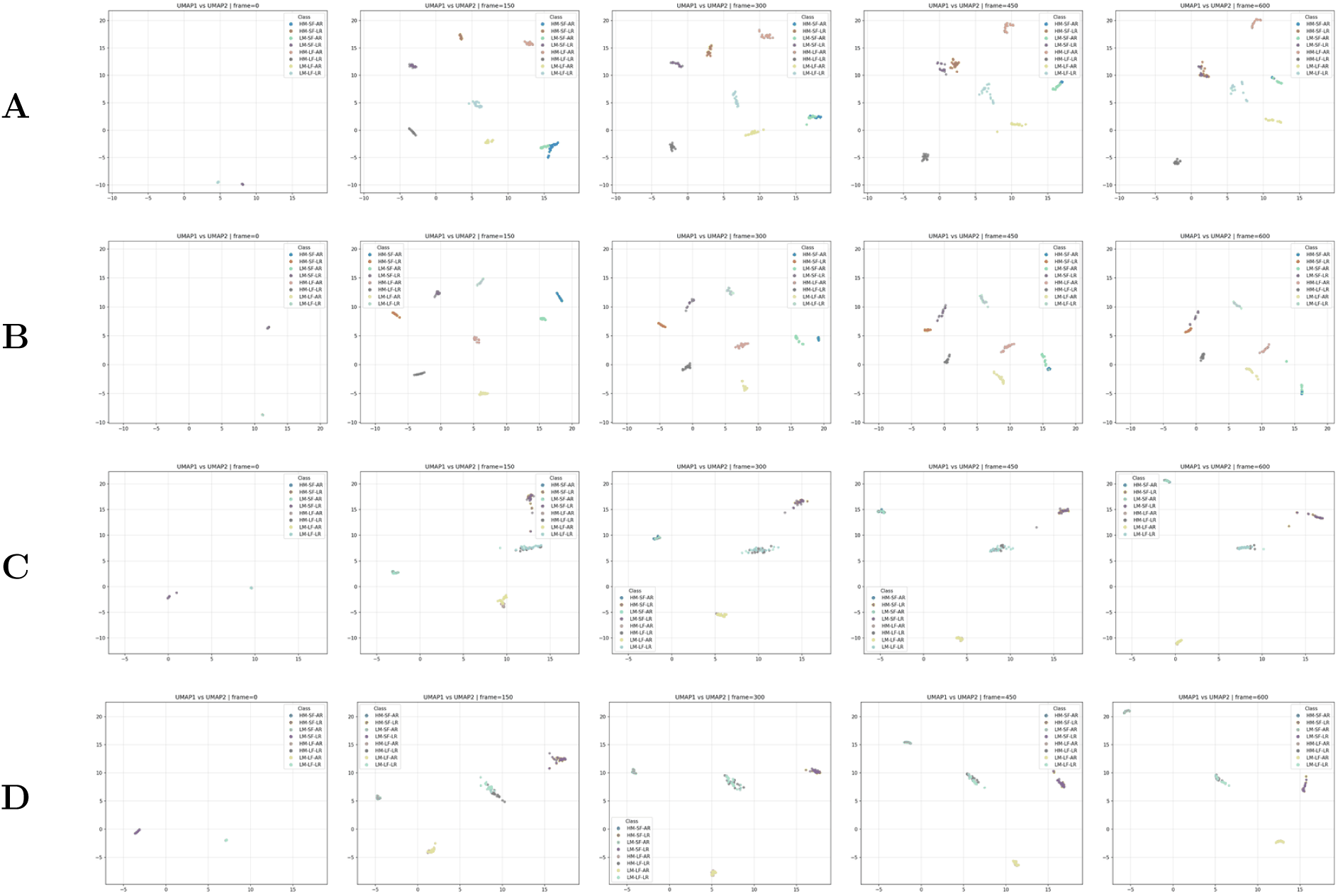
Evolution of trajectories of UMAP plots for the bins of the filament features for box of side=4 (A,C) and constrained cell of diameter=4 (B,D) including rigidities 0.075 pN µm*^−^*^1^ (AR) and 0.01 pN µm*^−^*^1^ (LR) for crosslinker densities 250 (A–B) and 15.625 (C–D). Columns: *(left to right)* frames=0, 150, 300, 450, 600.

These results show that both filament-feature PCA and UMAP can decode mechanical information encoded by our filament-level features, with temporal trajectories offering more insight.

#### 3.1.3 Haralick features capture global texture and collectively separate trajectories along the high motors-low motors axes

Haralick texture descriptors quantify spatial correlations in image intensity and capture global properties of network organization such as condensation, interstitial pore topology, and aggregate compactness. Because the fourteen Haralick features are highly correlated, there might be many redundant features. Dimensional reduction via PCA or UMAP is therefore necessary to explore a low-dimensional representation that captures dominant modes of texture variation while removing correlations and noise. We again look at the PCAs for changing motor density, filament length and filament rigidity together, looking at different PCAs for the different crosslinker densities. We now see though that the first 2 PCAs account for more than 85% of the explained variance for all the cases (Figure 10), allowing us to plot them against each other to gain insight about the systems as well. We can also use UMAP to present a lower dimensional space to visualize possible cluster formation in feature space.

**Figure 10:**
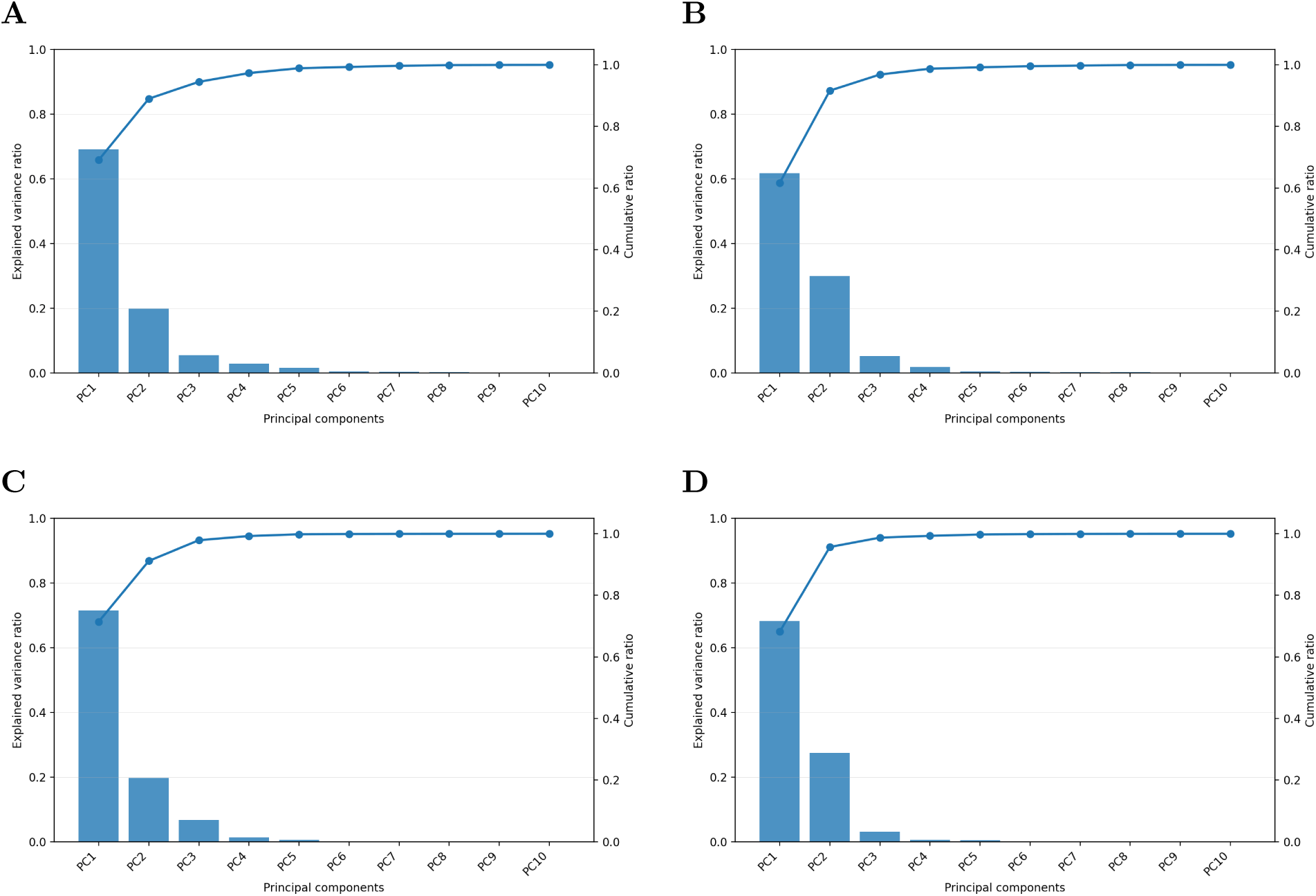
Explained variances for PCA plots for Haralick textural features along with time for box of side=4 (A,C) and constrained cell of diameter=4 (B,D) including rigidities 0.075 pN µm*^−^*^1^ (AR) and 0.01 pN µm*^−^*^1^ (LR) for crosslinker densities 250 (A-B) and 15.625 (C-D).

#### 3.1.4 Haralick-feature PCA and UMAP clusters trajectories primarily along the high and low motor axes and secondarily along the rigidity axes

Unlike for the filament-level measures, static PCA of Haralick features robustly separates low-motor from high-motor systems for low crosslinker density (Figure 11 Panel C-D). Low-motor systems cluster tightly due to weak reorganization of the texture, whereas the high-motor systems shift strongly in PCA space due to large-scale reorganization of textures. Similar results are seen when UMAPs are computed for the same standardized feature sets (Figure 12).

**Figure 11:**
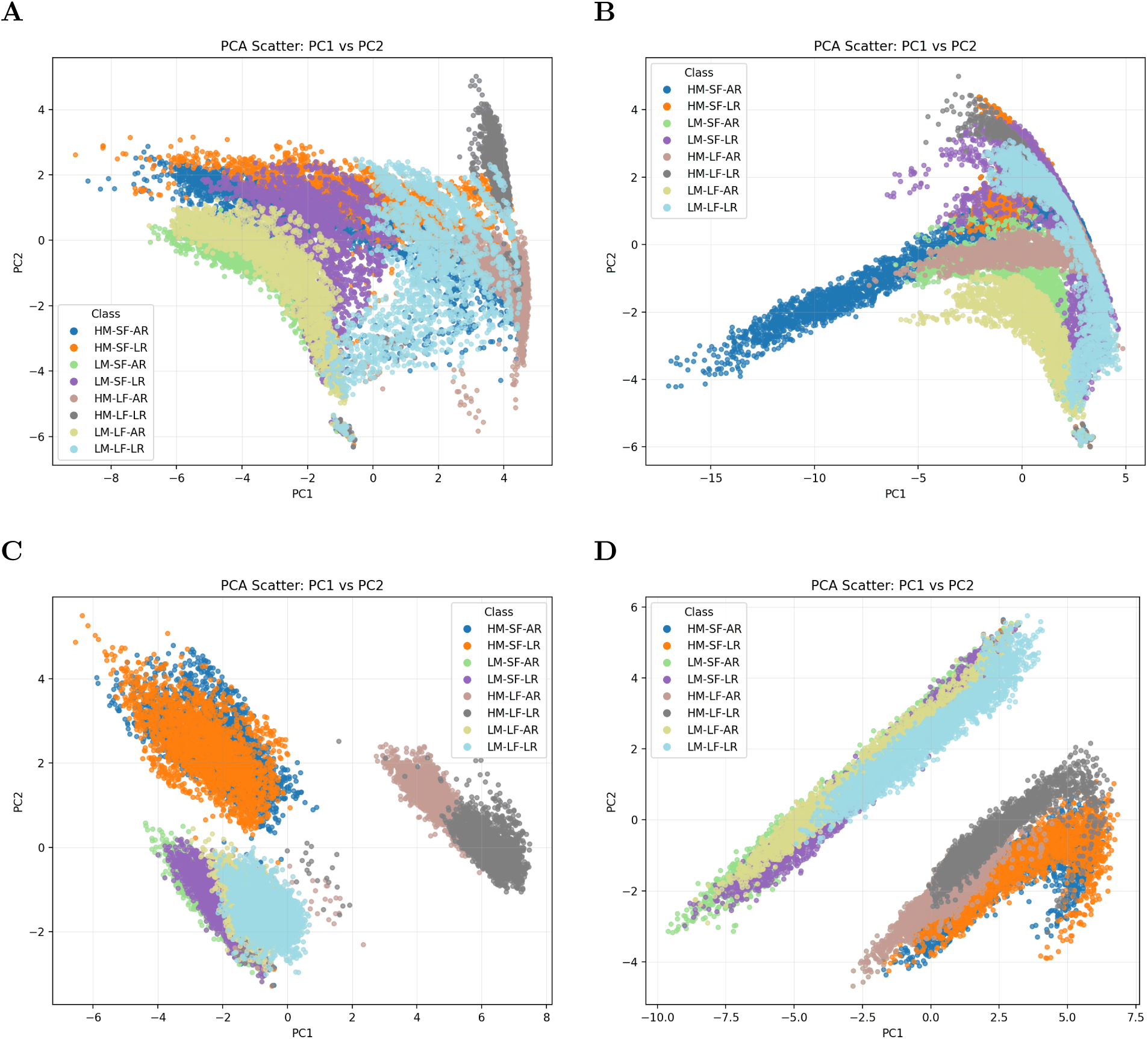
PCA plots for Haralick textural features along with time for box of side=4 (A,C) and constrained cell of diameter=4 (B,D) including rigidities 0.075 pN µm*^−^*^1^ (AR) and 0.01 pN µm*^−^*^1^ (LR) for crosslinker densities 250 (A-B) and 15.625 (C-D).

**Figure 12:**
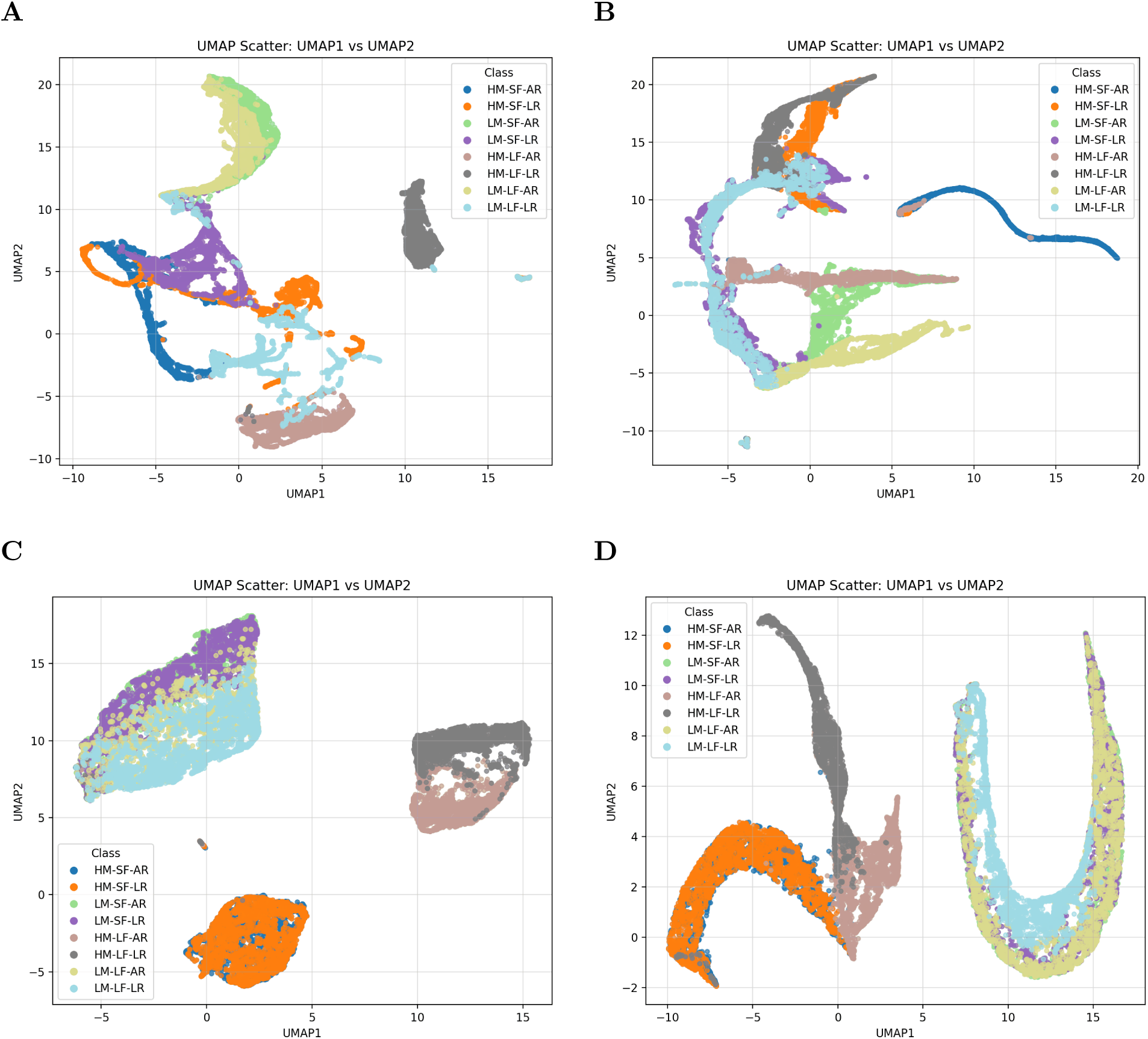
UMAP plots for Haralick textural features along with time for box of side=4 (A,C) and constrained cell of diameter=4 (B,D) including rigidities 0.075 pN µm*^−^*^1^ (AR) and 0.01 pN µm*^−^*^1^ (LR) for crosslinker densities 250 (A-B) and 15.625 (C-D).

At low crosslinker density, high-motor systems further separate by filament length in static Haralick PCA (Panel C), which is seemingly lost when looking at a circular constrained cell (Panel D). This is probably due to the fact that in the absence of sufficient bundling, there is a greater textural difference between a stalled network and clustered network (Figure 1, Panel III and IV). When constrained, the filaments come closer, allowing for more bundling, which reduces this textural differnce (Figure 2, Panel III and IV). Time-resolved Haralick PCA trajectories recover these transient distinctions (Figure 13).

**Figure 13:**
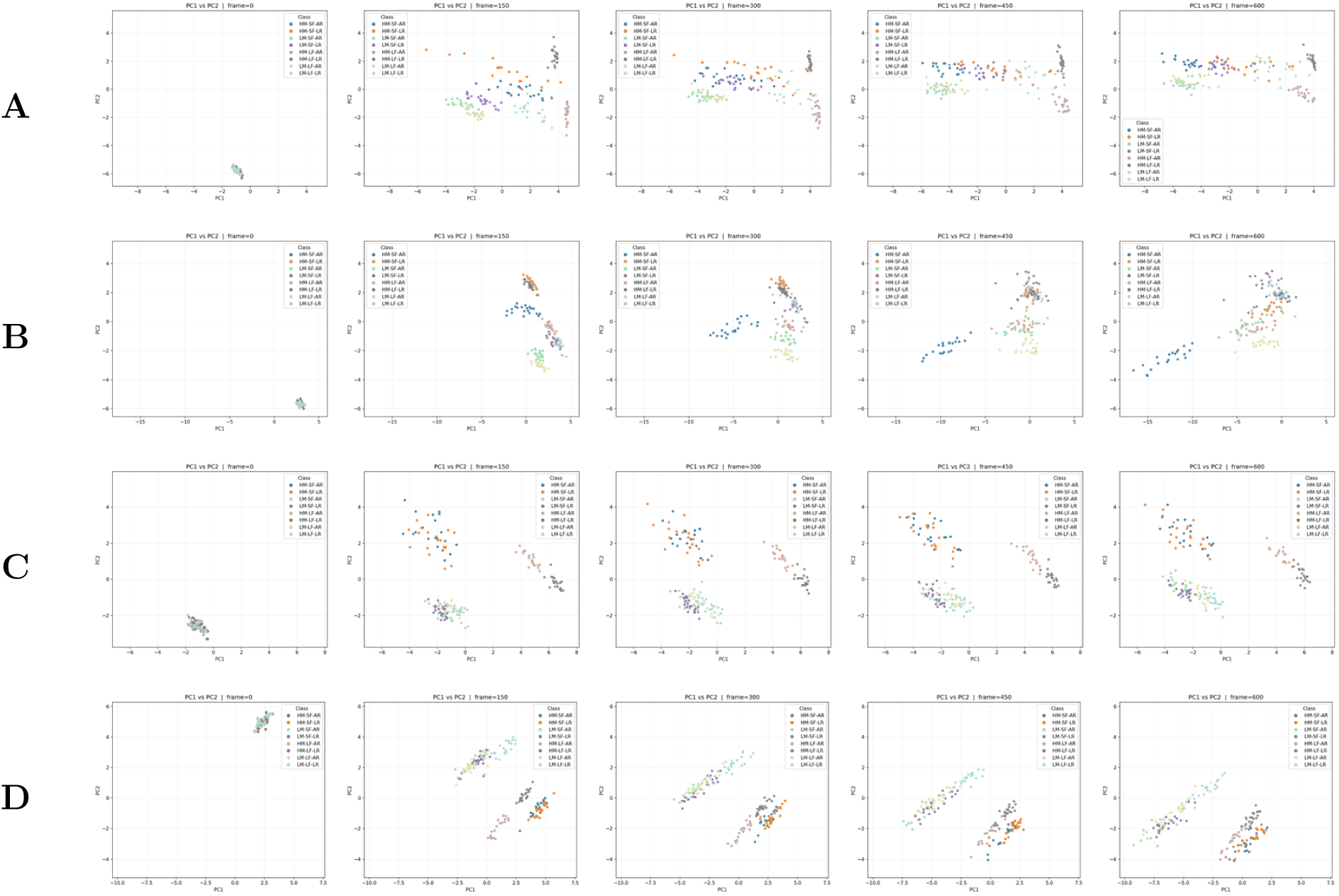
Evolution of trajectories of PCA plots for Haralick features for box of side=4 (A,C) and constrained cell of diameter=4 (B,D) including rigidities 0.075 pN µm*^−^*^1^ (AR) and 0.01 pN µm*^−^*^1^ (LR) for crosslinker densities 250 (A–B) and 15.625 (C–D). Columns: *(left to right)* frames=0, 150, 300, 450, 600.

**Figure 14:**
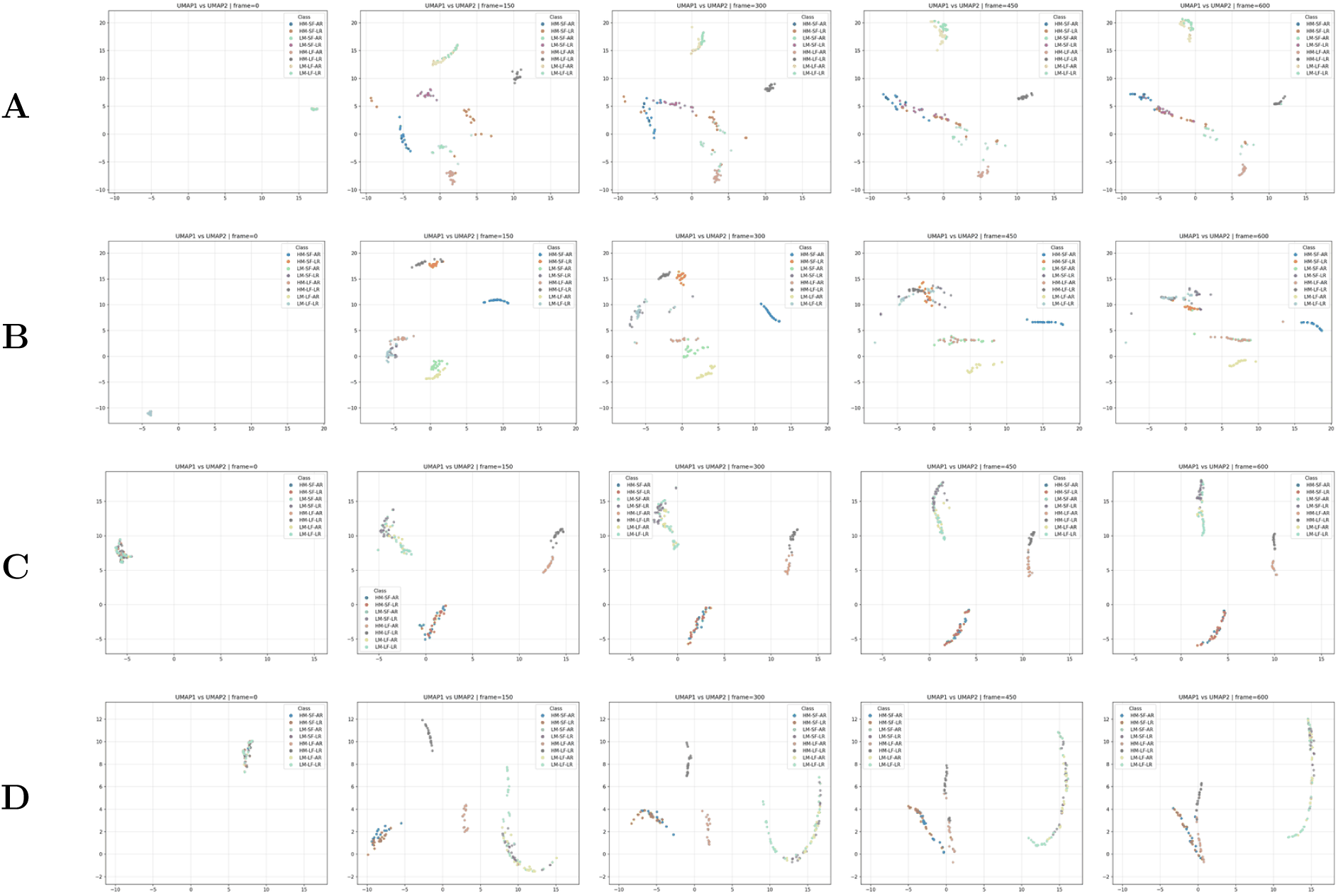
Evolution of trajectories of UMAP plots for Haralick features for box of side=4 (A,C) and constrained cell of diameter=4 (B,D) including rigidities 0.075 pN µm*^−^*^1^ (AR) and 0.01 pN µm*^−^*^1^ (LR) for crosslinker densities 250 (A–B) and 15.625 (C–D). Columns: *(left to right)* frames=0, 150, 300, 450, 600.

At high crosslinker density, this secondary separation is also reduced or lost from the static PCA plots (Panel A-B), as strong bundling probably produces similar coarse-grained textures at some point in the trajectories, despite underlying filament-scale differences. Looking at the time-resolved PCAs, while high-motor, short and long-filament systems follow distinct paths in texture space before converging once bundling saturates, the clustering by motor density is only good for low trajectory time, getting worse over time, as there is more mixing between clusters as trajectory progresses.

Looking at UMAP trajectories (Figure 2, there are some similar trends. For the low crosslinker cases (Panel C-D), the low motor cases also cluster together. But the high motor cases cluster separately for the two cases. For the box (Panel A), the high motor cases remain clustered in short and long filament groups, whereas for the constrained cell (Panel D), at low trajectory times, the short filament cases group together whereas the long filament groups cluster separately based on their rigidities, only slowly converging together as time progresses.

These results show that both PCA and UMAP of Haralick features can contain information about different motor regimes, but require analysis of their temporal trajectories to offer more insight.

#### 3.1.5 Complementary roles of filament and Haralick features

Together, these results demonstrate that filament and Haralick features capture complementary aspects of active network organization. Filament features resolve polymer-scale mechanics, rigidity effects, and filament-length-dependent bundling, while Haralick features robustly detect bundling and large-scale texture reorganization, especially emerging from differences in motor activity as well as intrinsic rigidity. Dimensionality reduction is informative in regimes with weak stabilization, whereas time-resolved trajectories are essential in regimes dominated by larger changes over the cases over time. Combining both feature families with trajectory-based analysis therefore provides a comprehensive framework for distinguishing active disordered systems across physical regimes.

### 3.2 Experimental validation: Haralick features distinguish cytoskeletal filament classes

While Haralick features have been used extensively for cell morphology studies, they have not yet to our knowledge been used to distinguish between different constituents of the cytoskeleton. Our analysis shows that Haralick features should be able to distinguish polymer networks of different rigidities from each other. To test this, we used a publicly available database of images of actin, microtubules and vimentin across substrates of increasing stiffness (see Materials and Methods). We calculated Haralick features at various Haralick distances for these images, and then performed a PCA, plotting the first 2 components and coloring them by filament classes (Figure 15).

**Figure 15:**
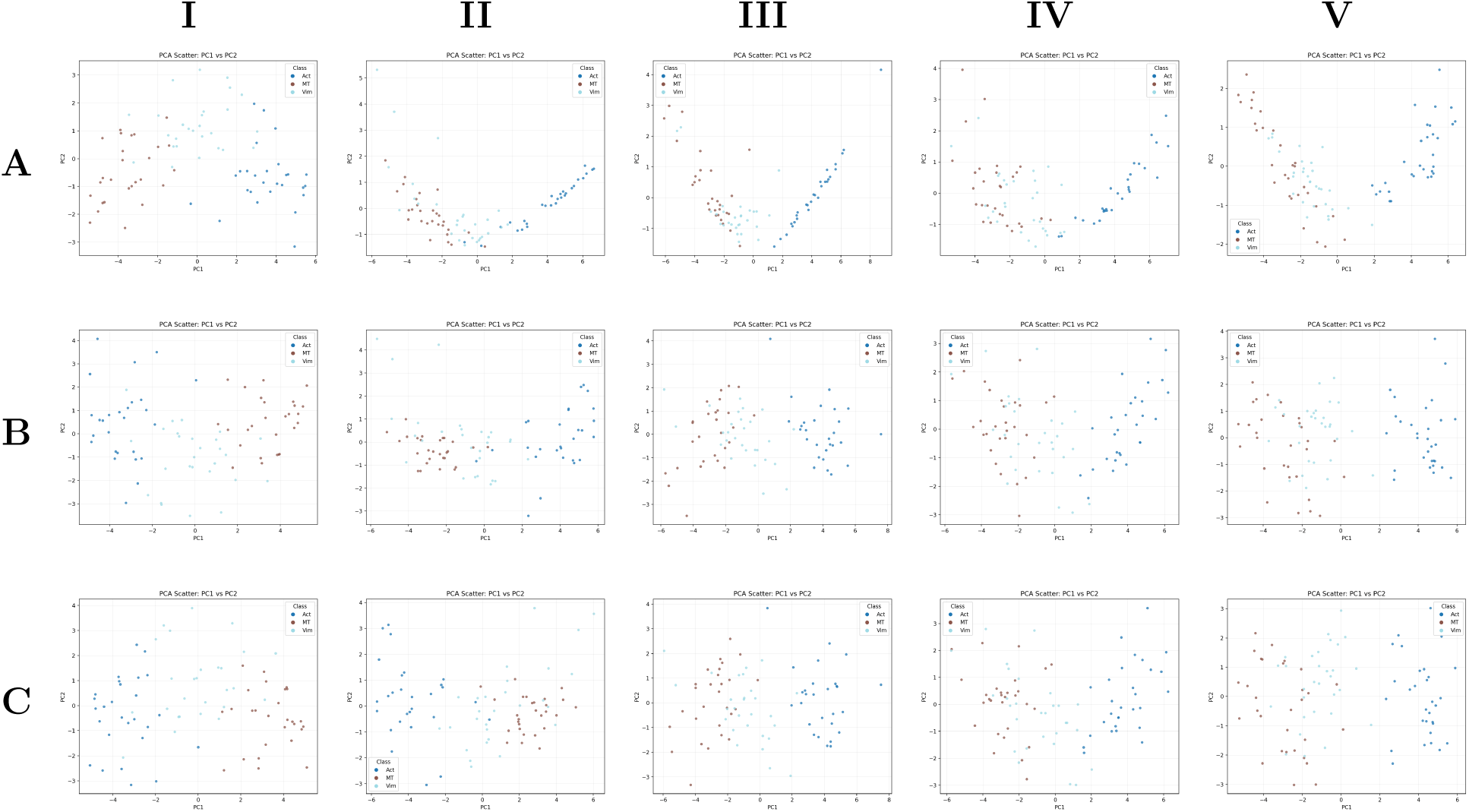
PCA of the haralick features for actin (Act), vimentin (Vim) and microtubules (MT) in experimental cells over substatrate with stiffness 1kPa (I), 5kPa (II), 10kPa (III), 30kPa (IV) and glass (V), calculated for haralick distance of 1 (A), 5 (B) and 10 (C)

Across all substrate stiffnesses and Haralick distances, actin consistently clusters separately from vimentin and microtubules. Vimentin and microtubules, in contrast, lie closer to each other in PCA space, showing more mixing for the cases where stiffness of the substrate is above 1*kPa* (Panel II-V).

Static PCA plots are sufficient to distinguish filament classes in the experimental data. Unlike the simulations, where systems evolve through condensation and bundling phases, the experimental images represent stabilized cellular states. As a result, temporal trajectories are not required to recover group structure.

These experimental results validate that texture-based morphospace analysis can discriminate between cytoskeletal systems with distinct mechanical properties in real biological contexts. Together with the simulation results, this demonstrates that Haralick features capture physically meaningful differences in filament organization.

## 4 Conclusion

In this paper we carried out simulations of a disordered polymer network in two dimensions, whose dynamics were determined by cross-linker and motor proteins, loosely based on actin filaments and myosin motors. For simplicity we did not consider polymerization processes, but assumed that filament length remains fixed. However, we altered filament rigidity and used two geometries for our simulations, a square box of size 4 *µm* and a circle with the same diameter, that increased the confinement of the polymers a little. We altered motor concentration and cross-linker concentration and studied the resulting 16 different cases using morphological features.

We concentrated on measures of filament bending (curvature and alignment order, referred to as filament features) and texture (Haralick features) since in our system the polymers did not show any nematic ordering and hence order parameters that were measuring this aspect of network organization were not very informative (see Supplementary Fig. SI1). We found that filament features did capture network dynamics involving bending and were successful in demarcating shorter and longer filaments due to this reason. To a lesser extent they could also distinguish between low motor and high motor cases especially during the more dynamic phase of network evolution. While no Haralick feature singly could distinguish between different trajectories at all times, collectively they demarcated clusters corresponding to differences in motor concentration as well as filament rigidities. Thus filament based features and textural features provide complementary information, that together were able to completely characterize the 16 separate cases that we studied here.

Since in simulations Haralick features captured differences in rigidity of the polymers, we decided to test them on a publicly available dataset that had images of actin, microtubules and vimentin. The results from the analysis of the experimental images show that actin Haralick features clustered separately, in 2d-PCA space, than vimentin and microtubules. These results highlight the usefulness of quantitative morphological descriptors as practical, interpretable tools for distinguishing cytoskeletal states in both simulations and real biological systems.

## Supplementary Information

**Figure SI1:**
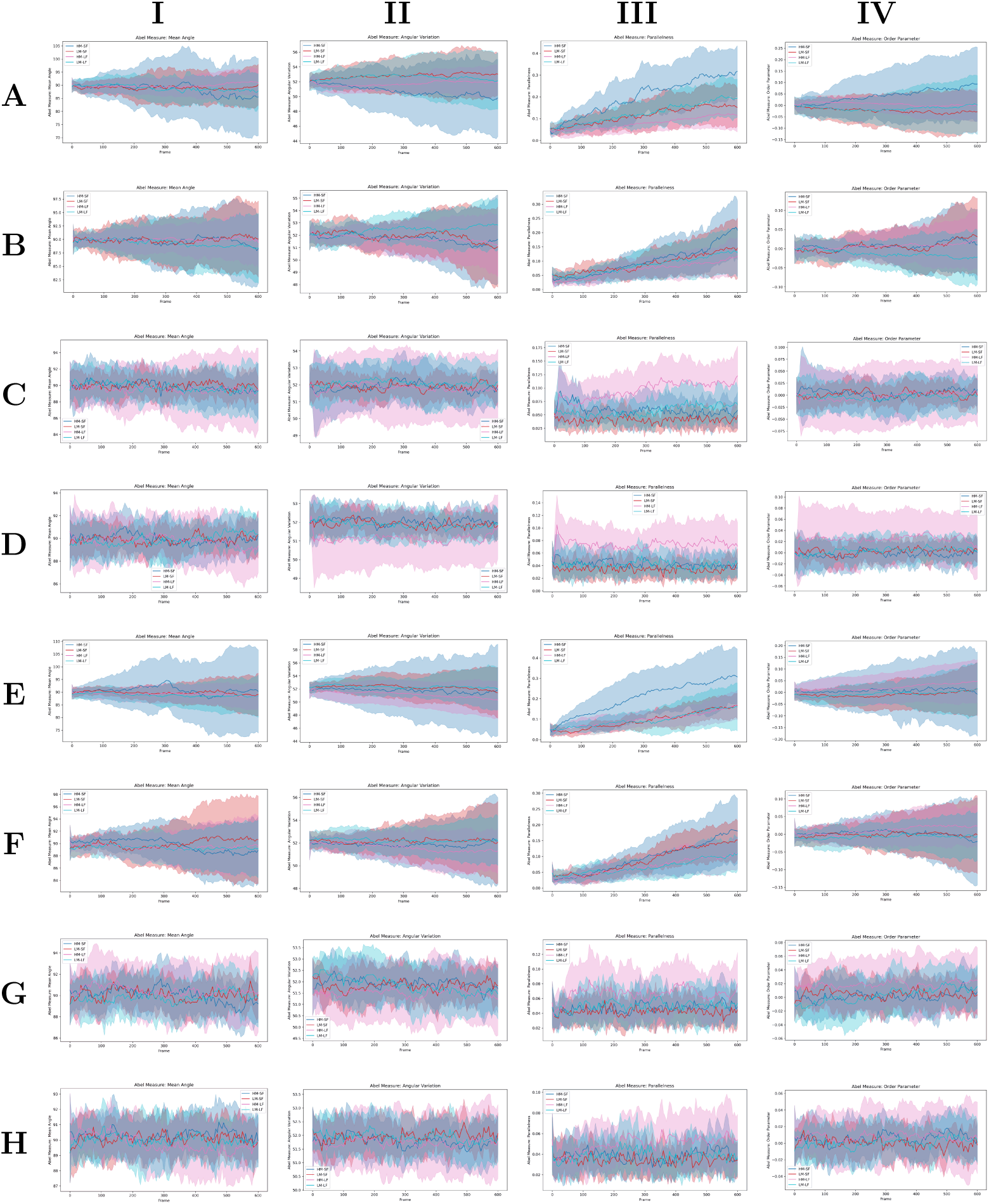
Plots for Mean Angle (I), Angular Variation (II), Parallelness (III) and Order Parameter (IV) for our simulation systems of box of side=4 (A-D) constrained cell of diameter=4 (E-H), having rigidities 0.075 pN µm*^−^*^1^ (A,C,E,G) and 0.01 pN µm*^−^*^1^ (B,D,F,H) for crosslinker densities 250 (A,B,E,F) and 15.625 (C,D,G,H).

**Figure SI2:**
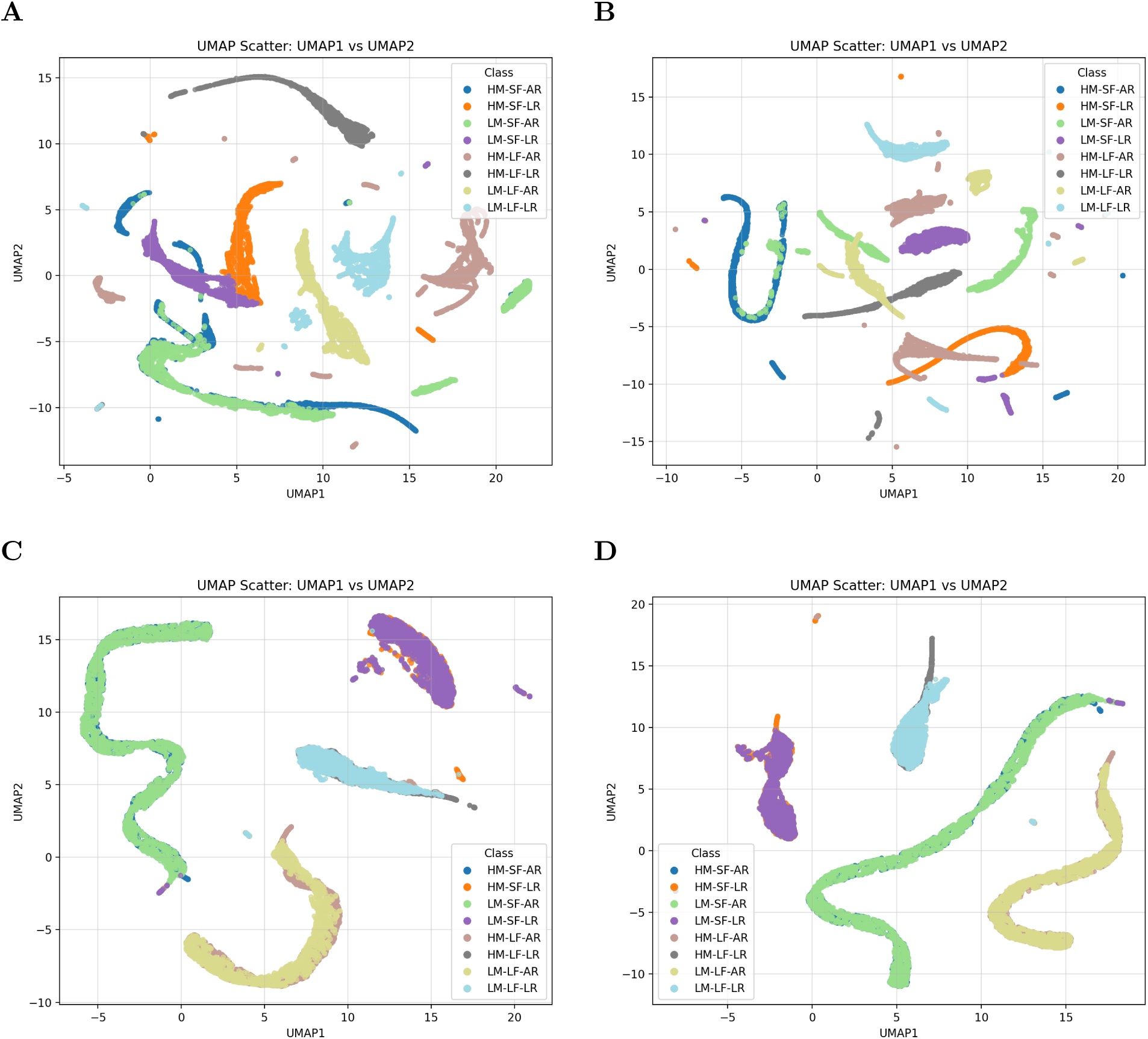
UMAP plots (with nearest neighbours=15) for the bins of the filament features along with time for box of side=4 (A,C) and constrained cell of diameter=4 (B,D) including rigidities 0.075 pN µm*^−^*^1^ (AR) and 0.01 pN µm*^−^*^1^ (LR) for crosslinker densities 250 (A-B) and 15.625 (C-D).

**Figure SI3:**
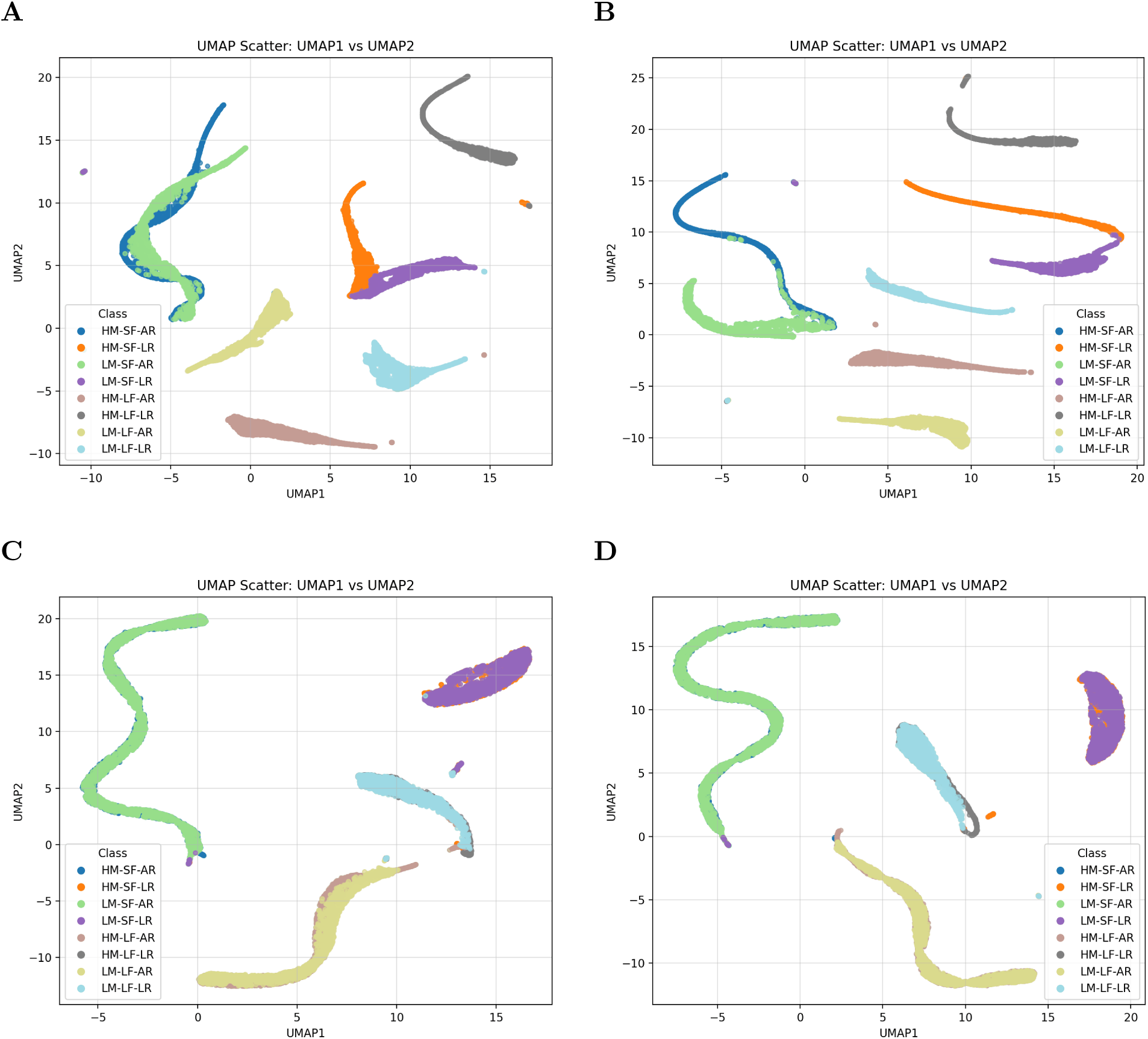
UMAP plots (with nearest neighbours=50) for the bins of the filament features along with time for box of side=4 (A,C) and constrained cell of diameter=4 (B,D) including rigidities 0.075 pN µm*^−^*^1^ (AR) and 0.01 pN µm*^−^*^1^ (LR) for crosslinker densities 250 (A-B) and 15.625 (C-D).

**Figure SI4:**
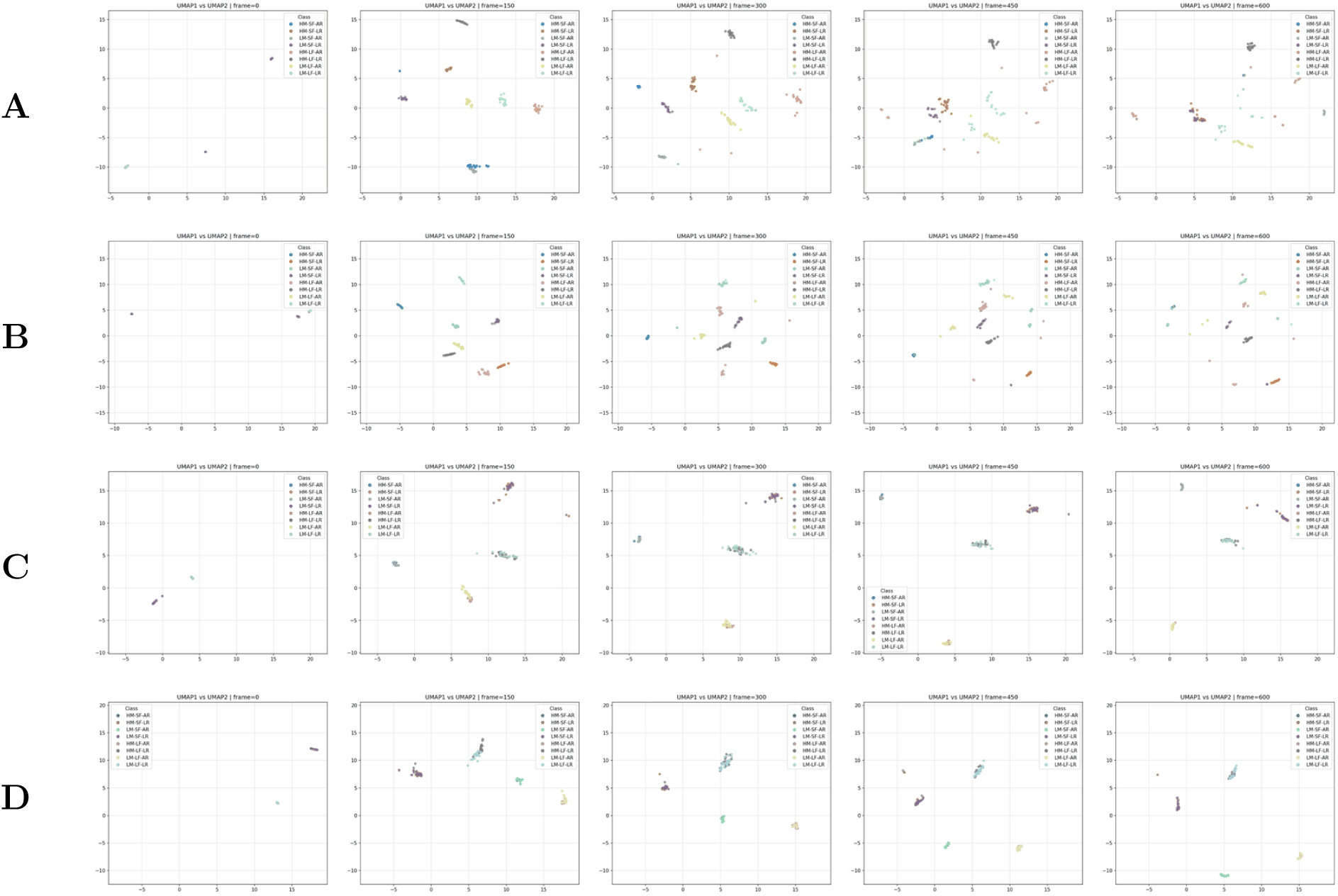
Evolution of trajectories of UMAP plots (with nearest neighbours=15) for the bins of the filament features for box of side=4 (A,C) and constrained cell of diameter=4 (B,D) including rigidities 0.075 pN µm*^−^*^1^ (AR) and 0.01 pN µm*^−^*^1^ (LR) for crosslinker densities 250 (A–B) and 15.625 (C–D). Columns: *(left to right)* frames=0, 150, 300, 450, 600.

**Figure SI5:**
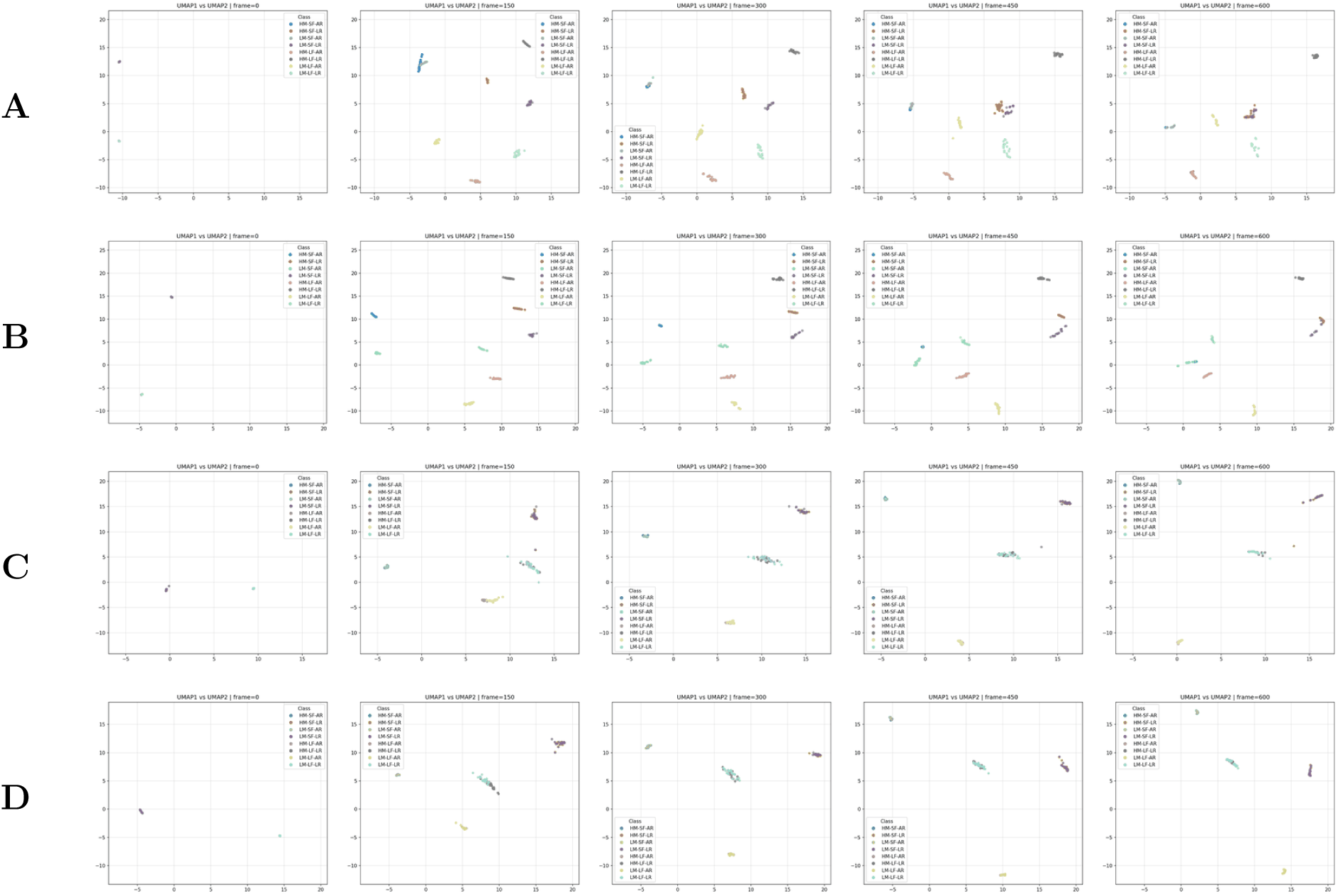
Evolution of trajectories of UMAP plots (with nearest neighbours=50) for the bins of the filament features for box of side=4 (A,C) and constrained cell of diameter=4 (B,D) including rigidities 0.075 pN µm*^−^*^1^ (AR) and 0.01 pN µm*^−^*^1^ (LR) for crosslinker densities 250 (A–B) and 15.625 (C–D). Columns: *(left to right)* frames=0, 150, 300, 450, 600.

**Figure SI6:**
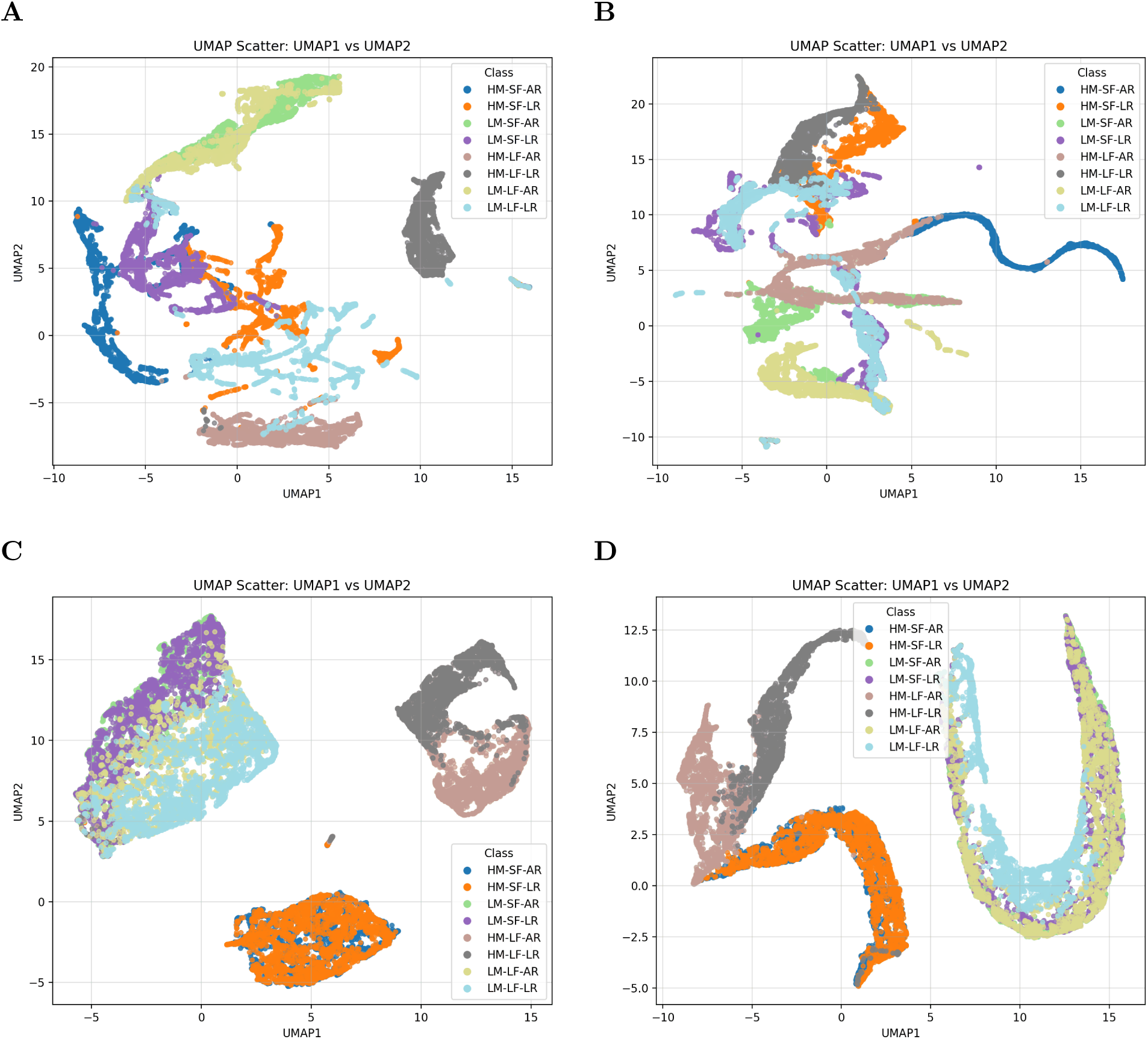
UMAP plots (with nearest neighbours=15) for Haralick textural features along with time for box of side=4 (A,C) and constrained cell of diameter=4 (B,D) including rigidities 0.075 pN µm*^−^*^1^ (AR) and 0.01 pN µm*^−^*^1^ (LR) for crosslinker densities 250 (A-B) and 15.625 (C-D).

**Figure SI7:**
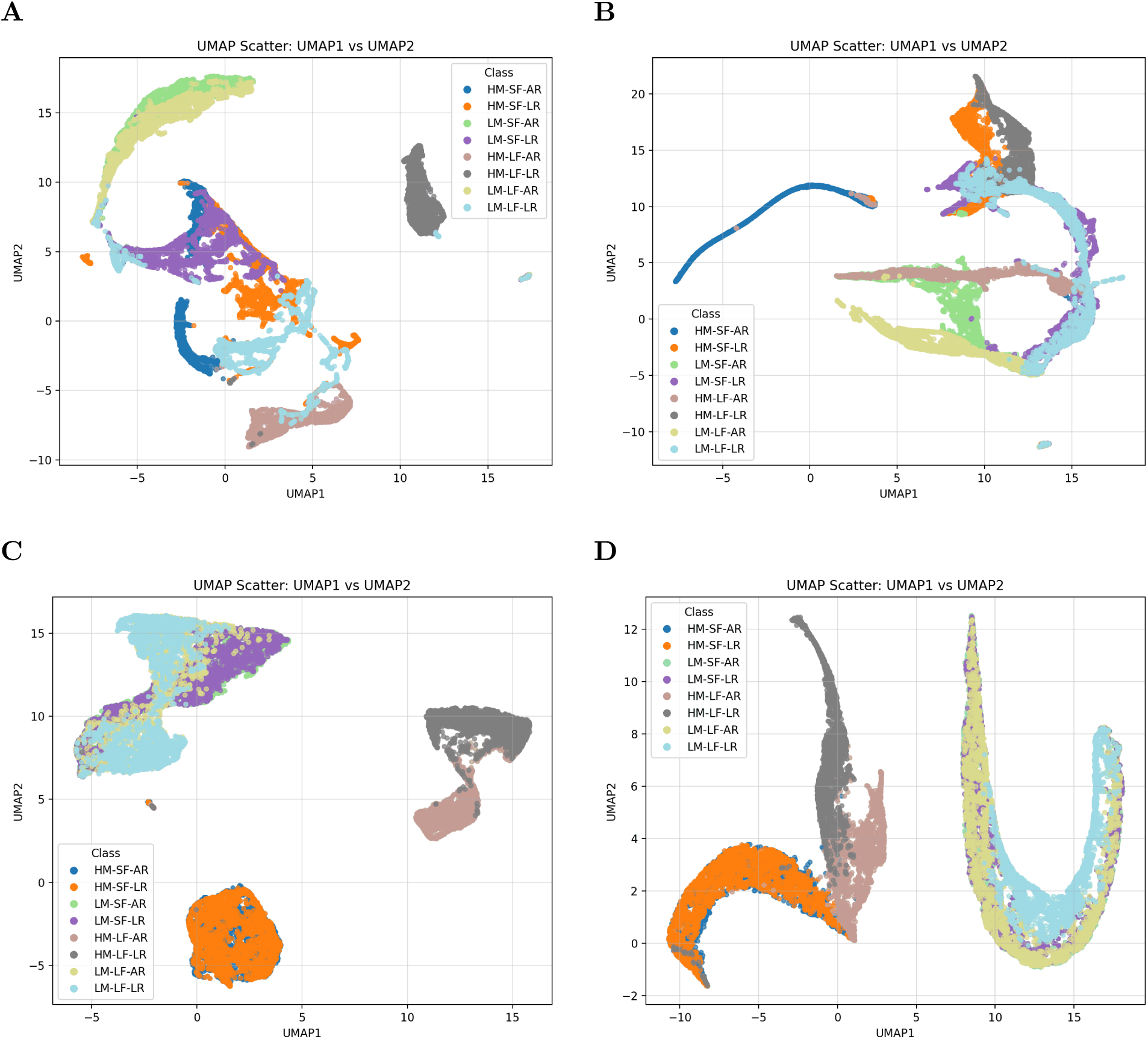
UMAP plots (with nearest neighbours=50) for Haralick textural features along with time for box of side=4 (A,C) and constrained cell of diameter=4 (B,D) including rigidities 0.075 pN µm*^−^*^1^ (AR) and 0.01 pN µm*^−^*^1^ (LR) for crosslinker densities 250 (A-B) and 15.625 (C-D).

**Figure SI8:**
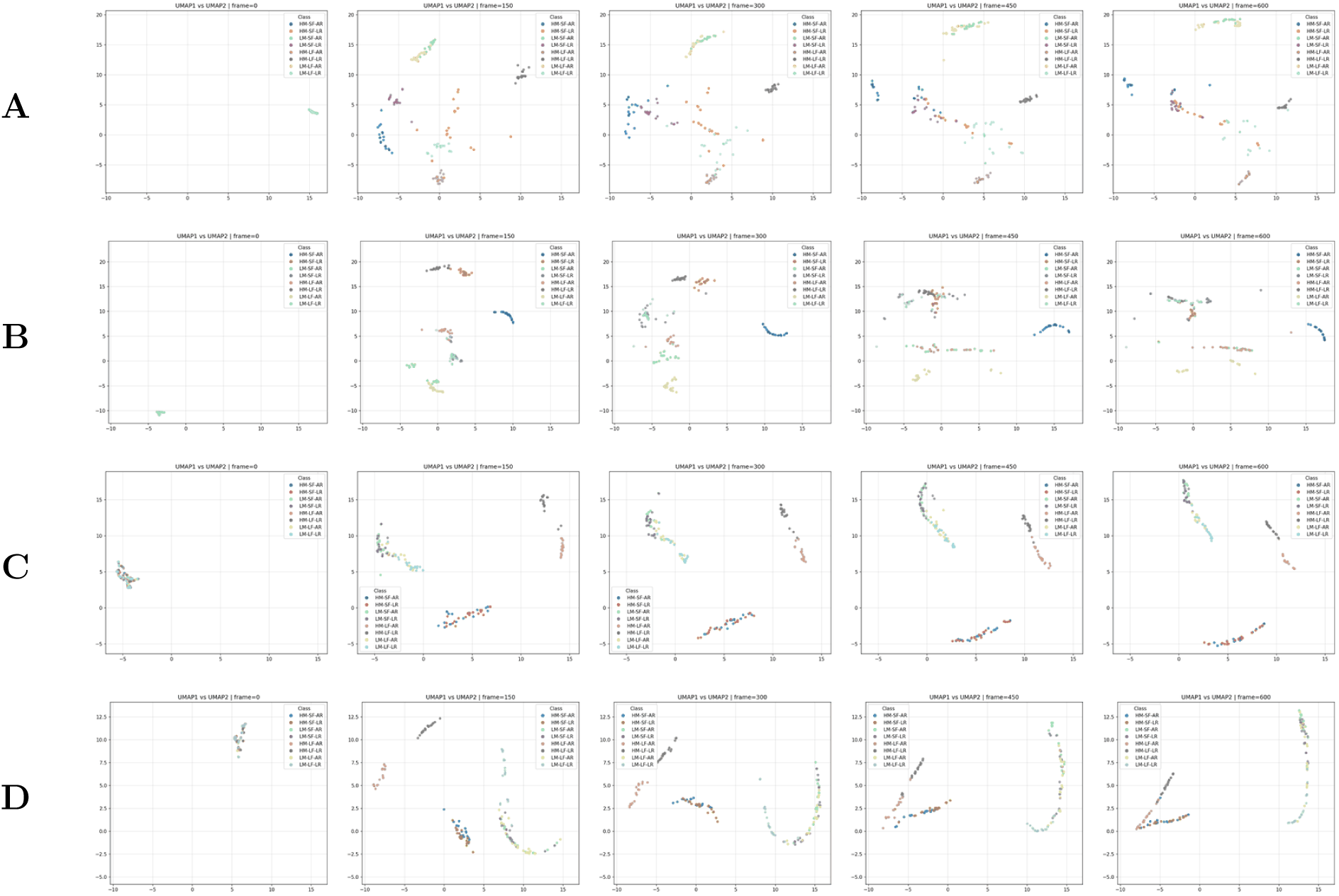
Evolution of trajectories of UMAP plots (with nearest neighbours=15) for Haralick features for box of side=4 (A,C) and constrained cell of diameter=4 (B,D) including rigidities 0.075 pN µm*^−^*^1^ (AR) and 0.01 pN µm*^−^*^1^ (LR) for crosslinker densities 250 (A–B) and 15.625 (C–D). Columns: *(left to right)* frames=0, 150, 300, 450, 600.

**Figure SI9:**
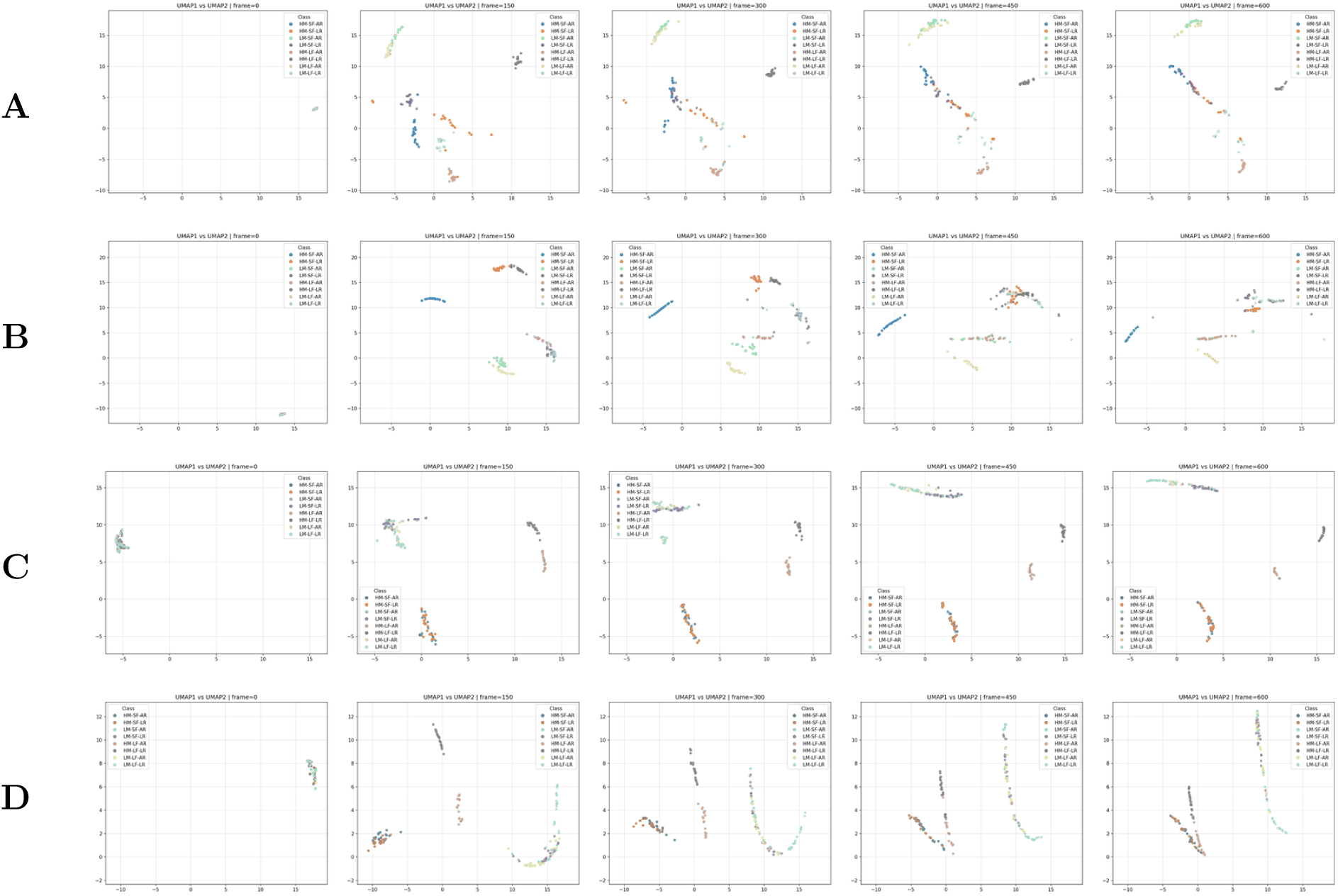
Evolution of trajectories of UMAP plots (with nearest neighbours=50) for Haralick features for box of side=4 (A,C) and constrained cell of diameter=4 (B,D) including rigidities 0.075 pN µm*^−^*^1^ (AR) and 0.01 pN µm*^−^*^1^ (LR) for crosslinker densities 250 (A–B) and 15.625 (C–D). Columns: *(left to right)* frames=0, 150, 300, 450, 600.

