## Supplementary Figures for "Morphological parameters can capture emergent properties of dynamic disordered cytoskeletal networks"

### Supplementary Information

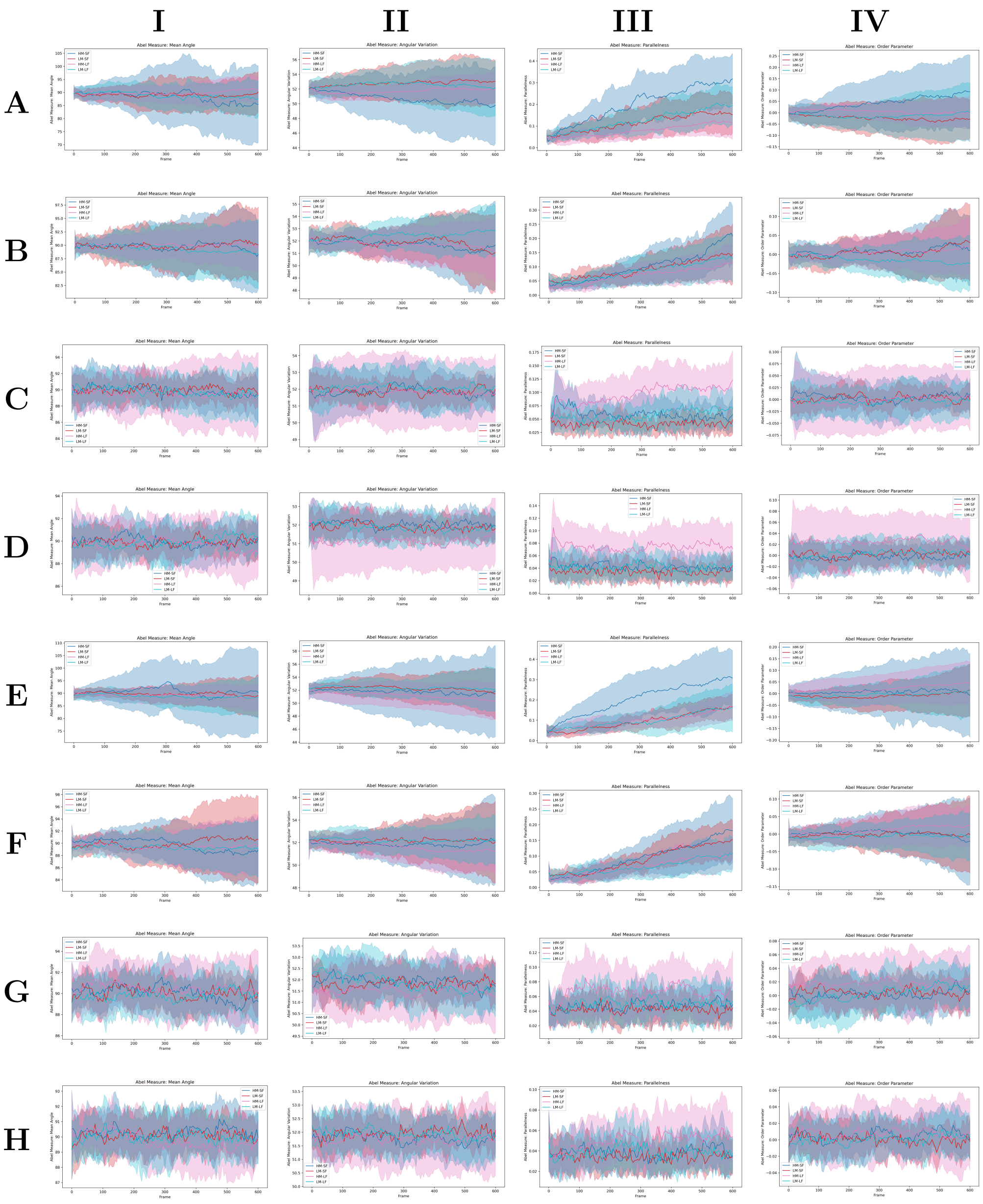


Figure SI1: Plots for Mean Angle (I), Angular Variation (II), Parallelness (III) and Order Parameter (IV) for our simulation systems of box of side=4 (A-D) constrained cell of diameter=4 (E-H), having rigidities 0.075 pN µm$^{-1}$ (A,C,E,G) and 0.01 pN µm$^{-1}$ (B,D,F,H) for crosslinker densities 250 (A,B,E,F) and 15.625 (C,D,G,H).

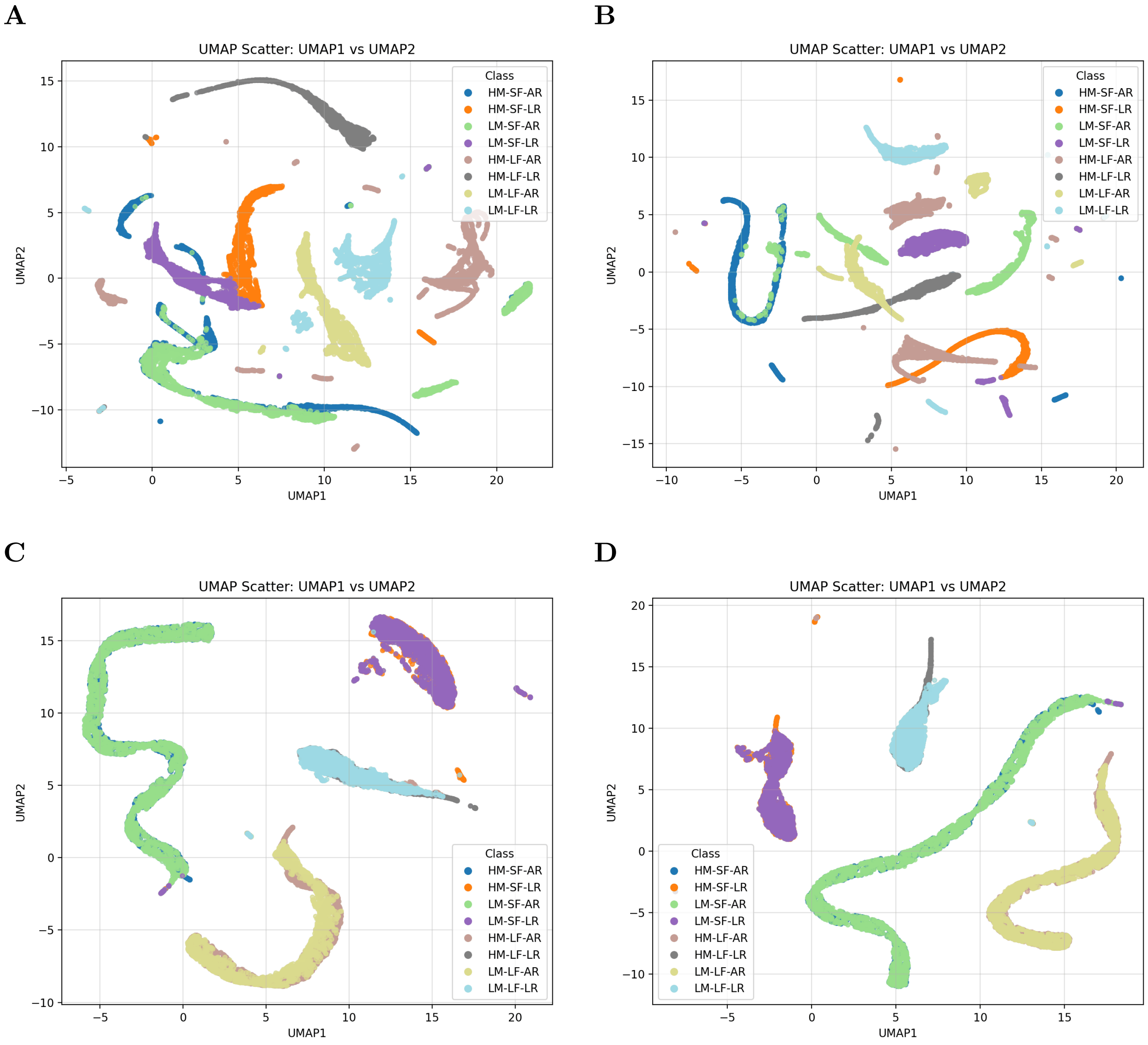


Figure SI2: UMAP plots (with nearest neighbours=15) for the bins of the filament features along with time for box of side=4 (A,C) and constrained cell of diameter=4 (B,D) including rigidities $0.075\,\mathrm{pN\,\mu m^{-1}}$ (AR) and $0.01\,\mathrm{pN\,\mu m^{-1}}$ (LR) for crosslinker densities 250 (A-B) and 15.625 (C-D).

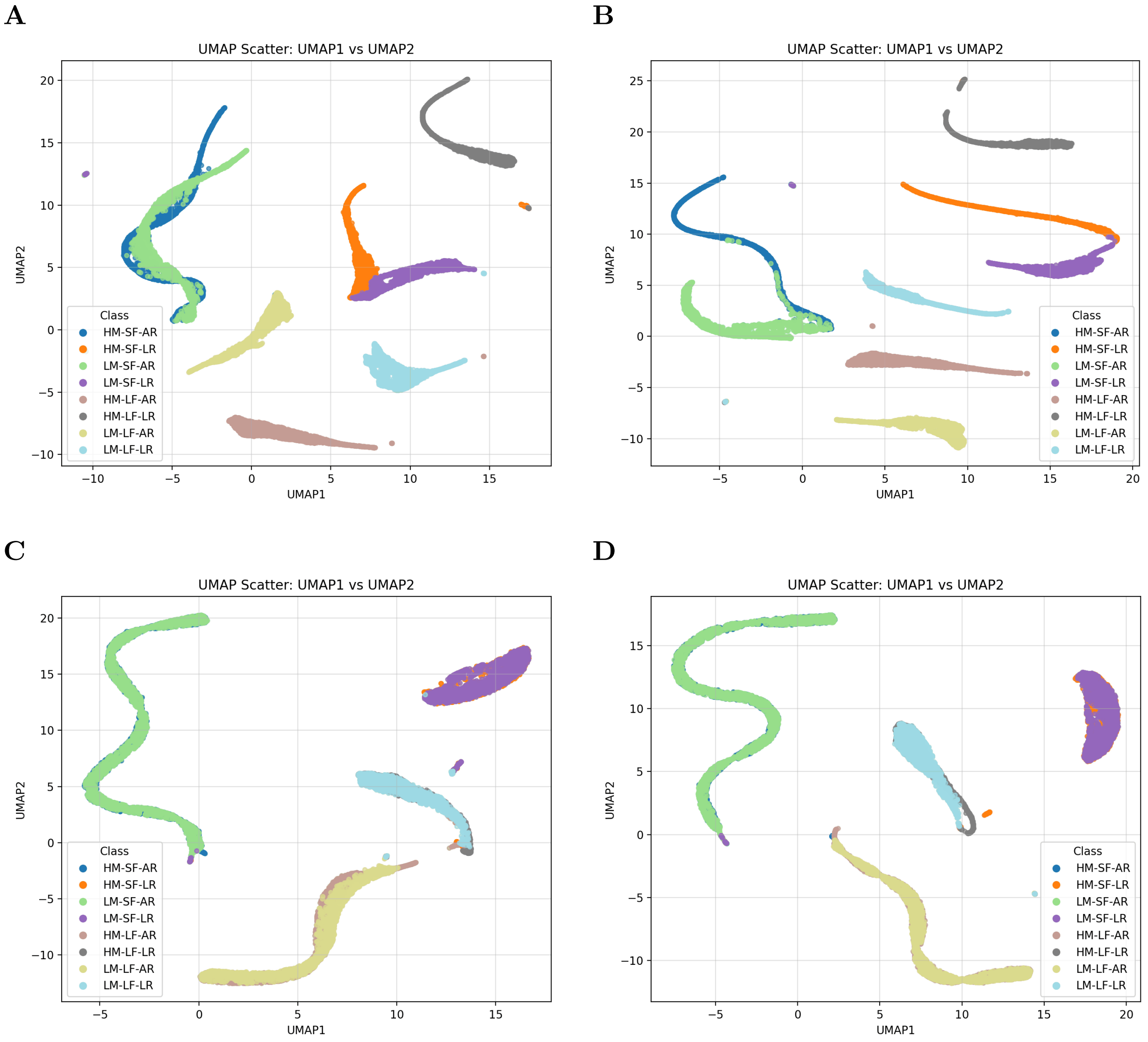


Figure SI3: UMAP plots (with nearest neighbours=50) for the bins of the filament features along with time for box of side=4 (A,C) and constrained cell of diameter=4 (B,D) including rigidities $0.075\,\mathrm{pN\,\mu m^{-1}}$ (AR) and $0.01\,\mathrm{pN\,\mu m^{-1}}$ (LR) for crosslinker densities 250 (A-B) and 15.625 (C-D).

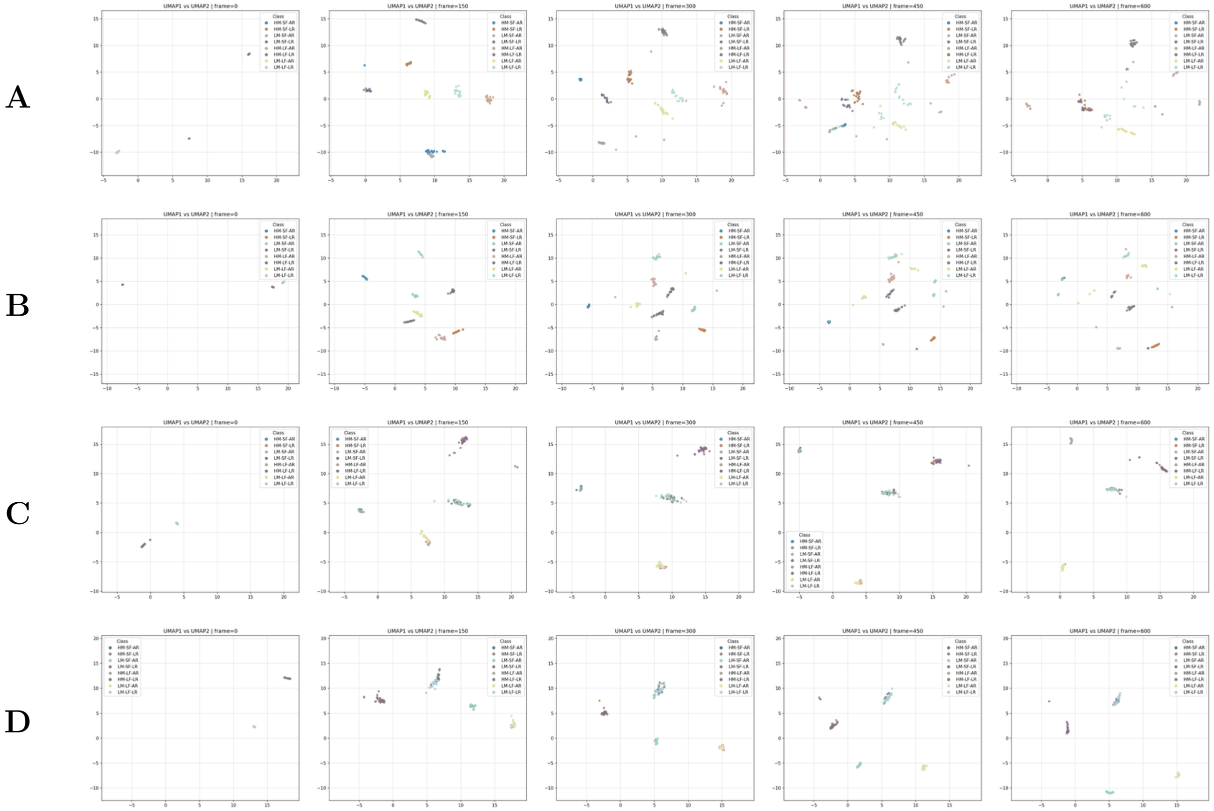


Figure SI4: Evolution of trajectories of UMAP plots (with nearest neighbours=15) for the bins of the filament features for box of side=4 (A,C) and constrained cell of diameter=4 (B,D) including rigidities $0.075\,\text{pN}\,\mu\text{m}^{-1}$ (AR) and $0.01\,\text{pN}\,\mu\text{m}^{-1}$ (LR) for crosslinker densities 250 (A–B) and 15.625 (C–D). Columns: *(left to right)* frames=$0, 150, 300, 450, 600$.

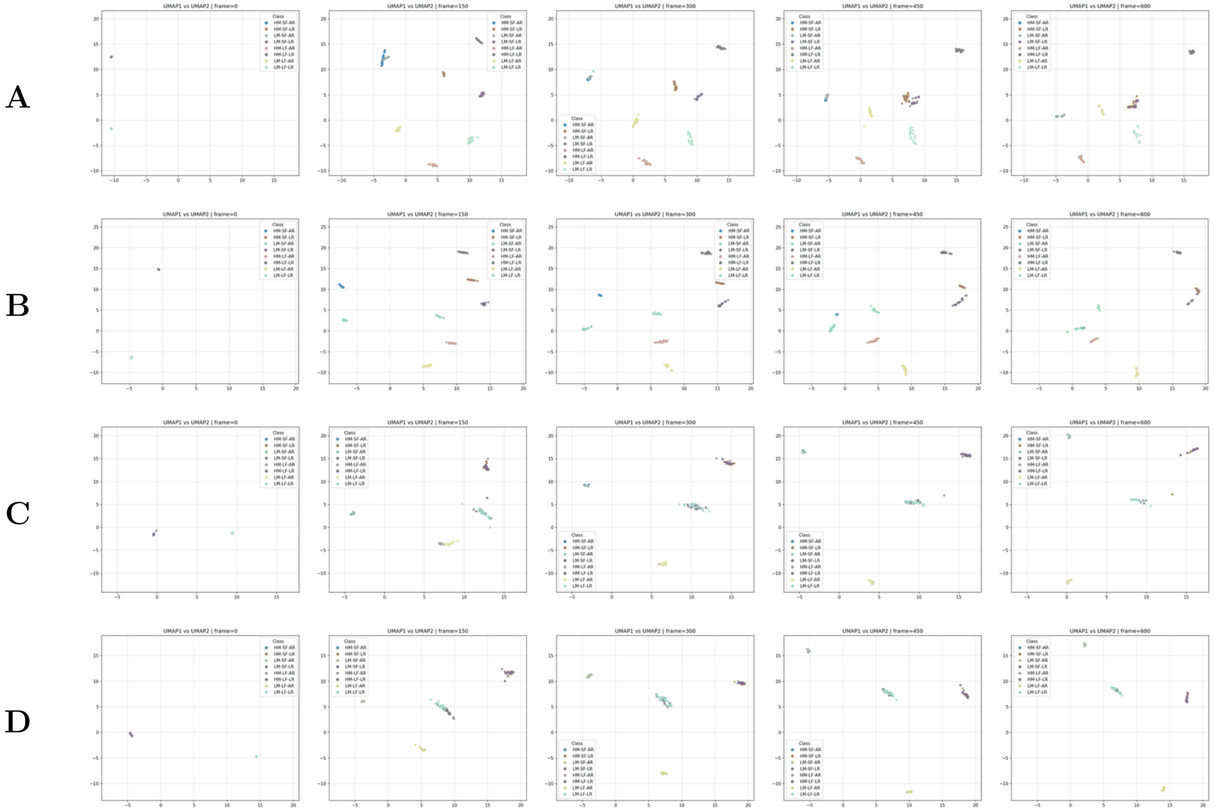


Figure SI5: Evolution of trajectories of UMAP plots (with nearest neighbours=50) for the bins of the filament features for box of side=4 (A,C) and constrained cell of diameter=4 (B,D) including rigidities $0.075\,\mathrm{pN\,\mu m^{-1}}$ (AR) and $0.01\,\mathrm{pN\,\mu m^{-1}}$ (LR) for crosslinker densities 250 (A–B) and 15.625 (C–D). Columns: *(left to right)* frames$=0, 150, 300, 450, 600$.

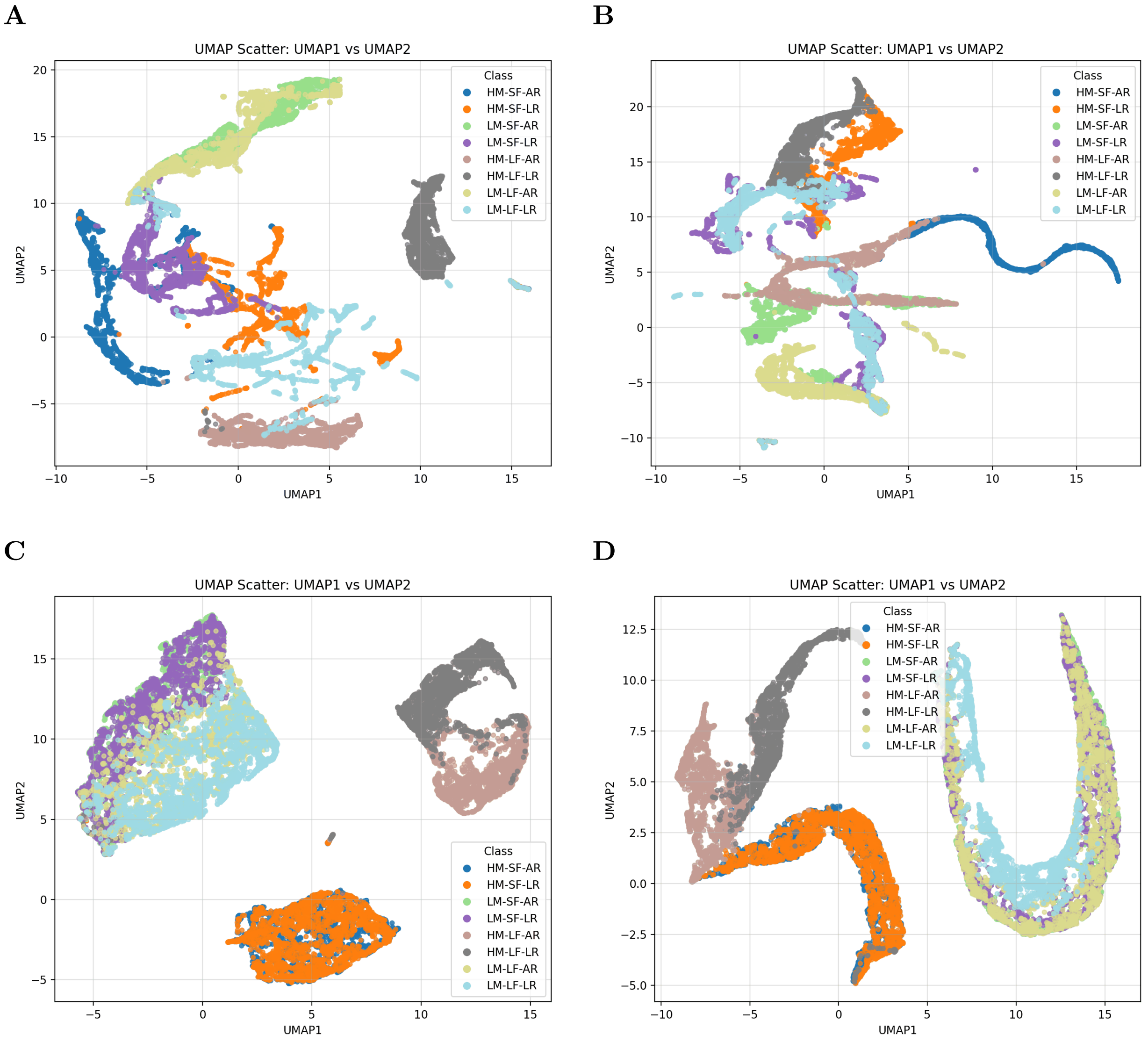


Figure SI6: UMAP plots (with nearest neighbours=15) for Haralick textural features along with time for box of side=4 (A,C) and constrained cell of diameter=4 (B,D) including rigidities $0.075\,\mathrm{pN\,\mu m^{-1}}$ (AR) and $0.01\,\mathrm{pN\,\mu m^{-1}}$ (LR) for crosslinker densities 250 (A-B) and 15.625 (C-D).

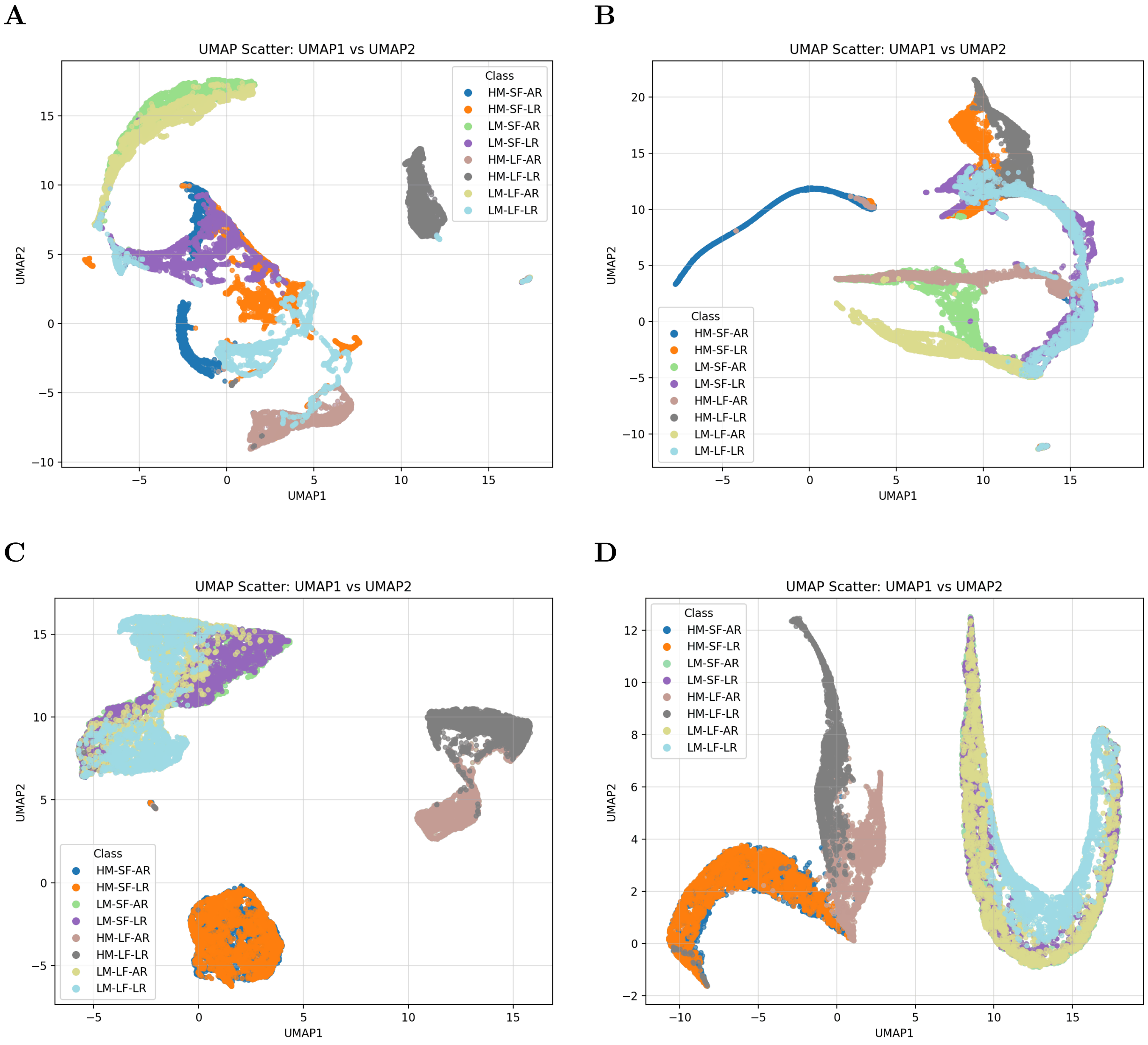


Figure SI7: UMAP plots (with nearest neighbours=50) for Haralick textural features along with time for box of side=4 (A,C) and constrained cell of diameter=4 (B,D) including rigidities $0.075\,\mathrm{pN\,\mu m^{-1}}$ (AR) and $0.01\,\mathrm{pN\,\mu m^{-1}}$ (LR) for crosslinker densities 250 (A-B) and 15.625 (C-D).

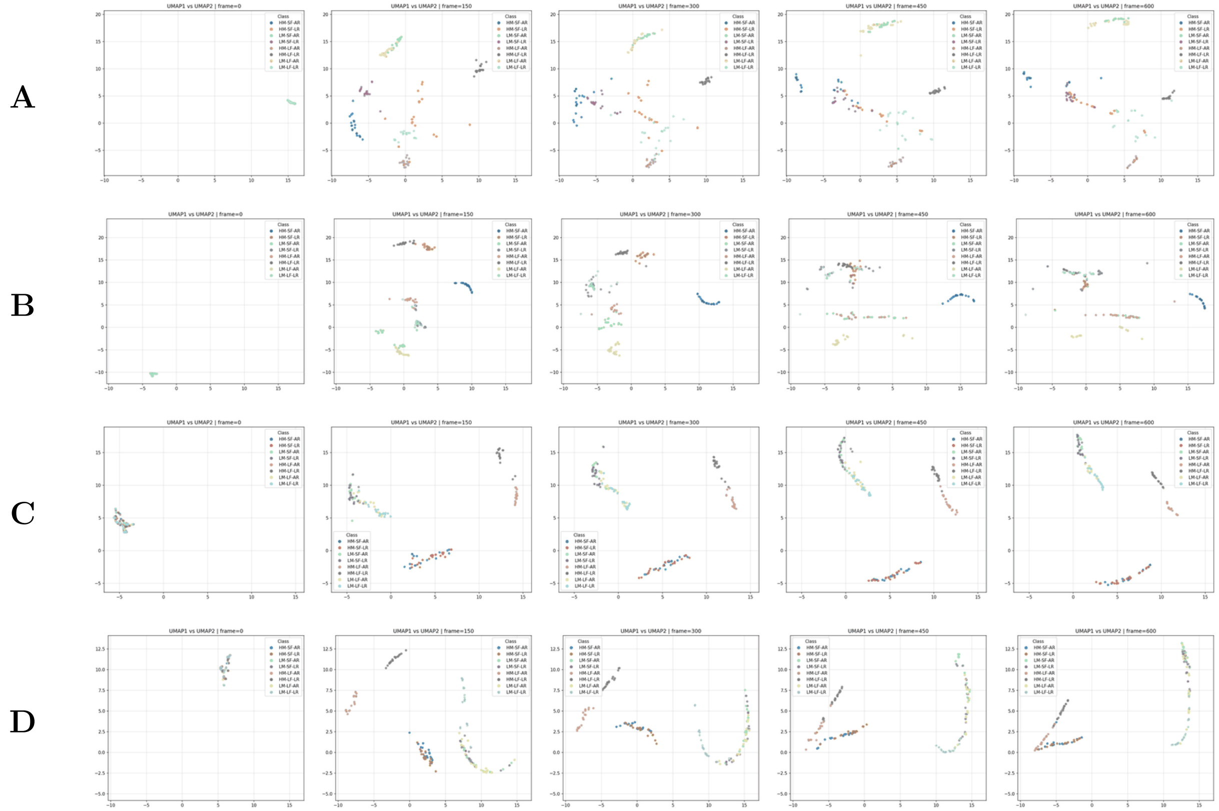


Figure SI8: Evolution of trajectories of UMAP plots (with nearest neighbours=15) for Haralick features for box of side=4 (A,C) and constrained cell of diameter=4 (B,D) including rigidities $0.075\,\mathrm{pN\,\mu m^{-1}}$ (AR) and $0.01\,\mathrm{pN\,\mu m^{-1}}$ (LR) for crosslinker densities 250 (A–B) and 15.625 (C–D). Columns: *(left to right)* frames$=0, 150, 300, 450, 600$.

A

B

C

D

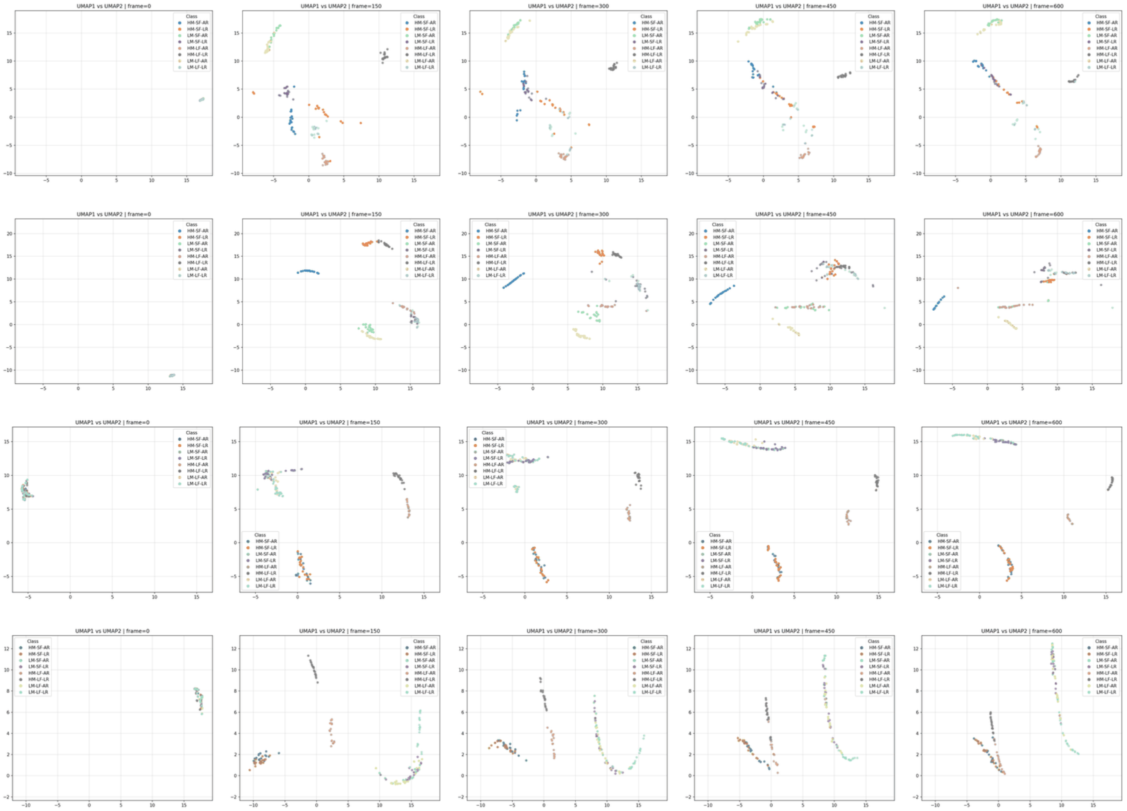


Figure SI9: Evolution of trajectories of UMAP plots (with nearest neighbours=50) for Haralick features for box of side=4 (A,C) and constrained cell of diameter=4 (B,D) including rigidities $0.075\,\mathrm{pN\,\mu m^{-1}}$ (AR) and $0.01\,\mathrm{pN\,\mu m^{-1}}$ (LR) for crosslinker densities 250 (A–B) and 15.625 (C–D). Columns: *(left to right)* frames=$0, 150, 300, 450, 600$.
